# The Annotated Blueprint: Integrated Functional Genomic Resources for a model Tetraploid Wheat *Triticum turgidum* cv. Kronos

**DOI:** 10.1101/2025.09.12.675711

**Authors:** Kyungyong Seong, Rakesh Kumar, Daniil M Prigozhin, China Lunde, Thales Henrique Cherubino Ribeiro, Sébastien Bélanger, Jo-Wei Allison Hsieh, McCree Tang, Blake C. Meyers, Ksenia V Krasileva

## Abstract

*Triticum turgidum* cv. Kronos is a tetraploid wheat cultivar that underpins one of the richest community platforms for functional genomics. Over the past decade, about 3,000 exome- and promoter-capture datasets, linked to mutagenized seed stocks, and transcriptomic and phenotypic resources have accumulated, yet the absence of a reference genome has constrained their impact. Here, we present a chromosome-scale reference genome of Kronos with high-confidence annotations, including manual curation of over 1,000 disease resistance (NLR) genes. This reference revealed previously hidden NLR diversity and clarified their genomic organization at chromosomal ends. Re-analysis of exome- and promoter-capture datasets enabled high-resolution mutation discovery in genes and regulatory regions that were previously inaccessible, uncovering the full standing variation present in Kronos mutant lines. We further re-curated transcriptomic and small RNA datasets, generating improved, genome-wide maps of microRNAs and phasiRNAs important for wheat development. Collectively, these resources elevate Kronos to reference quality and establish it as a versatile platform for functional and translational wheat research.

## Introduction

Wheat has sustained human civilization since its domestication over 10,000 years ago and is now cultivated on every continent except Antarctica (Salamini et al. 2002; Dubcovsky and Dvorak 2007; Shewry 2009; Levy and Feldman 2022). Through extensive breeding, wheat has been adapted to thrive across diverse climates, enabling year-round global harvests. Yet its success has been recently threatened by climate change, necessitating rapid germplasm adjustments and rational improvements (Mao et al. 2023; Alseekh et al. 2025). Driven by these challenges and its importance as a global staple, wheat has emerged as a tractable model system for scientific research (Krasileva et al. 2017; The International Wheat Genome Sequencing Consortium (IWGSC) et al. 2018; Adamski et al. 2020). Wheat grows readily in controlled environments, supports diverse genetic studies, and has highly plastic genomes that can withstand whole chromosome gains and losses as well as substantial mutational loads, making it well-suited for innovative experimental approaches (Bevan et al. 2017; Ghosh et al. 2018).

The adaptability of wheat is partly attributable to its allopolyploid nature (Dubcovsky and Dvorak 2007). Bread wheat (*Triticum aestivum*) is an allohexaploid (AABBDD), with a genome derived from three distinct progenitor species and a total size of ∼17 Gb (Marcussen et al. 2014). Its close relative, *Triticum turgidum*, is an allotetraploid (AABB), with a genome comprising ∼12 Gb (Maccaferri et al. 2019). Due to polyploidy, vast genome sizes, and extensive repeat content, generating high-quality wheat genomes has been a formidable challenge. Beginning in the early 2010s, international consortia launched coordinated efforts to overcome these obstacles by integrating up-to-date sequencing technology and algorithms, producing the first chromosome-level assembly of a wheat cultivar Chinese Spring (IWGSC et al. 2014; Zhu et al. 2021; Wang et al. 2025). More recently, advances in long-read sequencing technology, genome assembly algorithms, and scaffolding technology have enabled chromosome-scale assemblies from individual laboratories and accelerated pan-genomic analyses of wheat diversity (Avni et al. 2017; Maccaferri et al. 2019; Walkowiak et al. 2020; Athiyannan et al. 2022; Aury et al. 2022; Jiao et al. 2025) *T. turgidum* comprises diverse subspecies, commonly known as durum wheat, extensively cultivated for pasta production (Shewry 2009). One notable cultivar, *T. turgidum* cv. Kronos (hereafter referred to as Kronos), is a nonproprietary variety released in 1992 by Arizona Plant Breeders with Spring growth habit and suitability for late Fall planting in the Imperial and Central Valleys of California (https://goldenstategrains.com). Initially recognized for its excellent quality, Kronos has since become a foundational model system for basic and applied wheat research, widely adopted by academic laboratories and breeders worldwide. A major driver of this adoption is the development of EMS (ethyl methane sulphonate)-mutagenized TILLING (targeted induced local lesions in genomes) populations, which provide substantial genetic diversity (Uauy et al. 2009). For over 1,400 mutants, exome-capture (EC) and promoter-capture (PC) sequencing data have cataloged EMS-induced mutations across genic and promoter regions, enabling both forward and reverse genetic analyses (Krasileva et al. 2017; Zhang et al. 2023b).

Kronos mutant populations have been distributed through seed banks across three continents, supporting diverse functional studies. These encompass disease resistance (Wang et al. 2019; Lunde et al. 2025), yield-related traits (Jia et al. 2021; Niu et al. 2023), developmental processes (Shaw et al. 2019; Hawkins et al. 2021; Zhang et al. 2023a), and metabolic pathways (Wang et al. 2020b). Despite its widespread use, most functional analyses of Kronos have relied on the hexaploid bread wheat genomes, potentially obscuring variation and regulatory features that have uniquely arisen in tetraploid wheat since their divergence more than 10,000 years ago. Wheat genomes evolve rapidly, accumulating point mutations and small- and large-scale structural variations through recombination, transposition and duplication (Clavijo et al. 2017; IWGSC et al. 2018; Walkowiak et al. 2020; Jiao et al. 2025). A high-quality Kronos genome, together with its amenability for transformation and the availability of TILLING resources, fills a long-needed resource platform for forward and reverse genetic studies in *T. turgidum*.

The Kronos TILLING population is particularly valuable for studying disease resistance, a trait that requires continuous re-evaluation against evolving pathogens. Although Kronos initially showed moderate adult plant resistance to stripe rust and powdery mildew, it has become susceptible to these diseases, underscoring the need to identify and deploy new resistance (R) genes. Among R genes, nucleotide-binding leucine-rich repeat (NLR) receptors form a rapidly evolving and agronomically important gene family that mediates immunity (Jones and Dangl 2006; Jones et al. 2024). In hexaploid wheat, more than 3,400 NLR loci were identified, illustrating their remarkable diversity and complexity (Steuernagel et al. 2020). However, this diversity presents major challenges for accurate gene annotation, which is essential for continuous discovery and functional characterization against diverse fungal pathogens, such as *Puccinia graminis* (stem rust), *Puccinia striiformis* (stripe or yellow rust), *Magnaporthe oryzae* (wheat blast), *Zymoseptoria tritici* (Septoria blotch), and *Fusarium graminearum* (head blight), as well as bacterial pathogens like *Xanthomonas translucens* (bacterial leaf streak) (Dean et al. 2012; Singh et al. 2016; Sapkota et al. 2020). In most pangenome assemblies, NLRs are predicted through annotation transfer from reference genomes or approximated based on conserved motifs, without resolving full gene structures that could be critical for functional studies (Walkowiak et al. 2020; Jiao et al. 2025).

While advances in wheat genomics have focused mainly on protein-coding genes, long non-coding RNAs (lncRNAs) and small RNAs (sRNAs) also play pivotal roles along plant development. Among these, microRNAs (miRNAs) are produced from precursor transcripts encoded by microRNA genes (MIRs), the genomic loci that give rise to specific miRNA families. These miRNAs mediate post-transcriptional silencing of target mRNAs and lncRNAs, thereby regulating a wide array of biological processes (Borges and Martienssen 2015; Axtell and Meyers 2018). Most 21-nucleotide (nt) miRNAs induce cleavage of target RNAs, whereas 22-nt miRNAs can trigger biogenesis of secondary siRNAs, known as phased small interfering RNAs (phasiRNAs) (Liu et al. 2020). Loci encoding phasiRNA precursor transcripts are referred to as *PHAS* loci and may originate from either protein-coding genes or lncRNA-producing non-coding regions. Coding gene-derived *PHAS* loci stem from diverse gene families and produce 21-nt phasiRNAs involved in both vegetative and reproductive developments (Liu et al. 2020). By contrast, lncRNA-derived *PHAS* loci produce both 21-nt and 24-nt phasiRNAs, especially abundant in reproductive tissues. In grasses, 21-nt phasiRNAs predominantly accumulate at the premeiotic stage, functioning in both vegetative and reproductive tissues (Zhai et al. 2015; Fei et al. 2016; Liu et al. 2020; Bélanger et al. 2025), while 24-nt phasiRNAs peak at premeiotic or meiotic stages and appear to be specific to male reproductive development (Bélanger et al. 2020, 2024; Zhan et al. 2024).

In this study, we present an integrated suite of functional genomic resources for both basic and applied wheat research using *T. turgidum* cv. Kronos (Fig. 1). We generated a chromosome-scale genome assembly using PacBio long-read sequencing and scaffolding using chromosome conformation capture (Hi-C) data, resolving seven chromosomes from telomere to telomere and anchoring the remaining seven with one telomeric end. Evidence-based annotations achieved 99.9% BUSCO (Benchmarking Universal Single-Copy Orthologs) completeness, providing a high-confidence gene set. To advance disease resistance studies, we manually curated over 2,300 NLR loci, identifying 1,089 reliable NLRs, and quantified their sequence diversity at the pan-genome level. We also re-curated *PHAS* loci, revealing improved chromosome-level distribution patterns. About 3,000 exome capture and promoter capture datasets were remapped to the Kronos genome, enabling high-resolution mutation detection in genic and regulatory regions. Finally, we reanalyzed transcriptomic datasets, uncovering constitutive expression of NLRs in shoot apical meristem development. Collectively, these integrated resources establish Kronos as a reference-quality system for forward and reverse genetics, accelerating efforts to dissect gene function and enhance resilience in polyploid wheat (Table S1).

**Figure 1.**
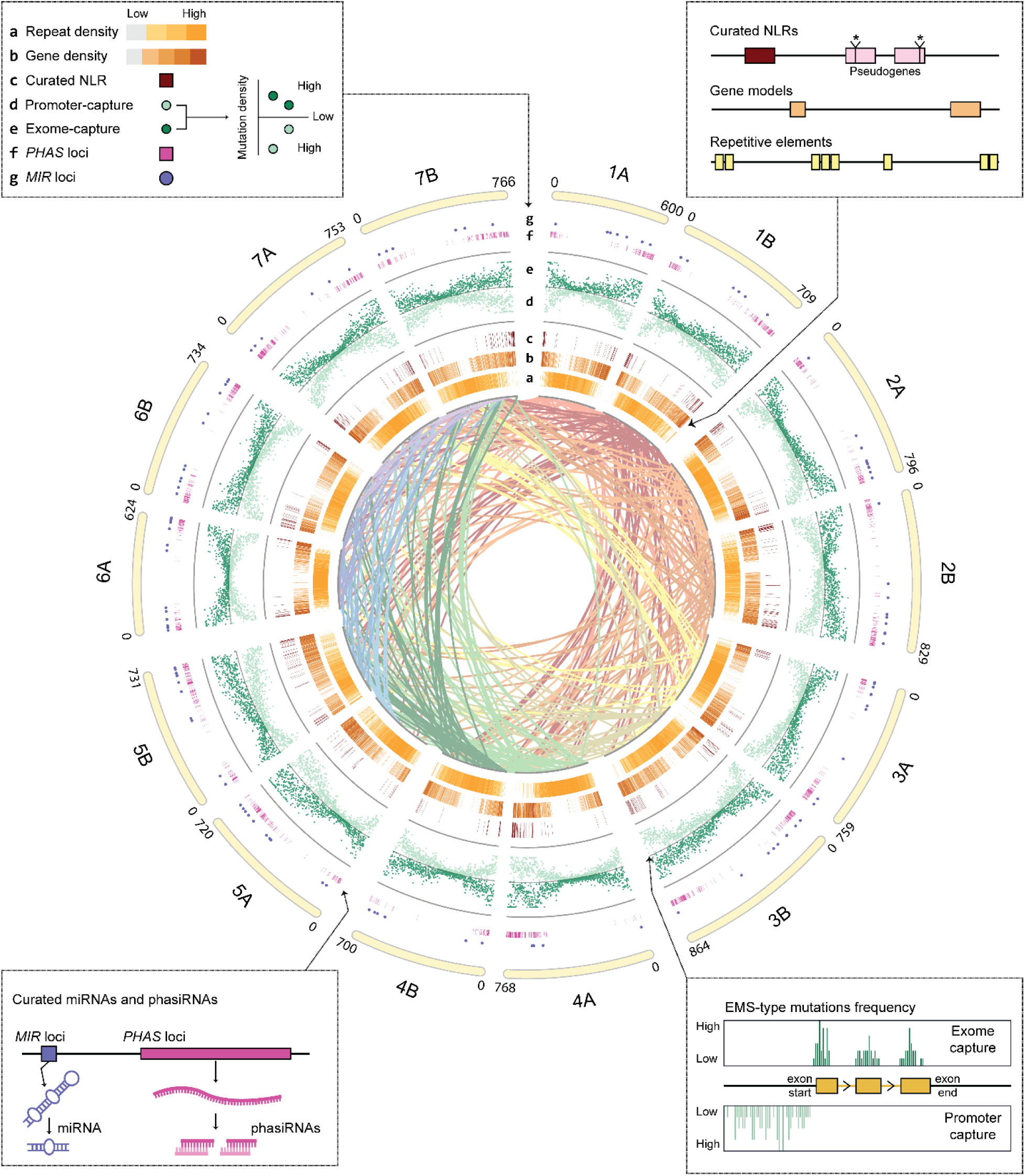
Integrated functional genomic resources for *Triticum turgidum* cv Kronos. Circos plot displaying the 14 Kronos chromosomes annotated with multi-layered genomic features (Krzywinski et al. 2009). Central ribbons indicate cross-chromosomal synteny relationships (>150 kb). Outer tracks display: (a) repeat density; (b) gene density; (c) high- and medium-confidence manually curated nucleotide-binding leucine-rich repeat (NLR) genes; (d and e) EMS-induced mutation frequency from promoter-capture and exome-capture sequencing, respectively; (f) *PHAS* loci encoding phased small interfering RNAs (phasiRNAs); and (g) *MIR* loci encoding microRNA (miRNAs). Chromosome lengths (in Mb) are shown in the outermost ring. Insets highlight key resources generated in this study.

## Results

### A chromosome-scale reference genome for Kronos with refined annotation

*T. turgidum* rarely outcrosses and maintains a high level of homozygosity through self-pollination. K-mer profiling of long-read whole-genome sequencing data for Kronos estimated a genome size of 10.45 Gb with 90.5% homozygosity, similar to another cultivated *T. turgidum* variety Svevo (Fig. S1; Table S2) (Maccaferri et al. 2019; Ranallo-Benavidez et al. 2020). Therefore, we generated a haplotype-collapsed (AB) assembly, consistent with the strategy used for Svevo. Using 50x high-fidelity (HiFi) reads, the initial assembly reached N50 of 41 Mb (Table S3). Hi-C scaffolding anchored 98.0% of the assembled sequence by length, reconstructing all 14 chromosomes totaling 10.35 Gb (Figs S2 and S3; Table S4). Telomeric repeats were detected on every chromosome, with seven fully spanning telomere-to-telomere, supporting near-complete assembly (Fig. S4). The remaining 210 Mb of unplaced contigs were scaffolded into a single sequence designated ‘Un’.

Our v1.0 protein-coding gene annotation integrated public short-read transcriptomic data available for Kronos and contained 69,808 high-confidence genes with 98.9% BUSCO completeness (Table S5). To further refine untranslated regions and correct mis-annotations, we incorporated public long-read transcriptomic data for other *Triticum* species, yielding v2.0 with 71,357 genes and 99.9% BUSCO completeness (Table S5). Orthology assessment with OMArk placed 70,630 (99.0%) genes into taxonomically consistent orthologous groups (Fig. S5; Table S6) (Nevers et al. 2025). In comparison, currently available Svevo annotation reports 66,559 genes with 97.0% BUSCO completeness (Table S5). To evaluate potential gene count inflation in our annotations, we applied the gene selection strategy used for Svevo (≥90% bidirectional coverage with UniProt homologs) (Maccaferri et al. 2019). This filtering retained 70,382 Kronos genes (Table S7). Given the marginal impact, we kept the full v2.0 annotations.

### Curated Kronos NLR repertoire distinguishes reliable genes from pseudogenes

NLR disease resistance genes are notoriously difficult to annotate automatically due to frequent duplication, clustered organization, and rapid diversification through both point mutation and structural variation, including birth and death processes (Barragan and Weigel 2021). We therefore manually curated the Kronos NLR repertoire, annotating 2,328 loci (Table S8). In an initial round of curation, we assessed their multi-domain architecture and homology to sequences in the NCBI non-redundant databases (Figs. S6-S8). In a subsequent iteration, we assigned confidence levels to each gene, using domain structure, homology, exon–intron integrity, expression support, and local duplication context (Table S8). Of the 2,328 loci, 1,239 (53%) were classified as low-confidence pseudogenes, typically comprising fragmented nucleotide-binding (NB-ARC) domains or truncated exons disrupted by splice-site mutations, frameshifts, or premature stop codons. The remaining 1,028 high- and 61 medium-confidence genes were designated reliable and used for downstream analyses (Table S8).

We evaluated representation of these reliable NLRs in the Kronos reference annotations. Of 1,089 NLRs, 755 (69%) in v1.0 and 846 (78%) in v2.0 had identical protein sequences to curated models (Table S9). For external validation of completeness, we compared against NLR-Annotator, which marks putative NLR loci but does not provide gene structures (Steuernagel et al. 2020). All loci detected by NLR-Annotator yet missing in our final annotations were pseudogenes, whereas 53 genes were missed by NLR-Annotator (Table S8). We integrated the 1,089 reliable NLR genes into annotation v2.1 and removed models at low-confidence loci, further improving the overall quality of the Kronos genome annotation (Table S5).

### Kronos NLR repertoire includes orthologs of cloned resistance genes

In hexaploid wheat, NLRs are enriched near distal chromosome arms, where elevated recombination and mutation can accelerate sequence diversification (Steuernagel et al. 2020). Consistent with previous findings, the distribution of NLRs in Kronos resembled that of Chinese Spring (Fig. 2A). NLR counts per chromosome were also comparable between the two genomes (Table S10) (Steuernagel et al. 2020). To explore the functional relevance of the Kronos NLR repertoire, we surveyed for orthologs of cloned NLRs conferring resistance to fungal diseases (Table S11). Putative Kronos orthologs were identified for leaf rust resistance genes *Lr10*, *Rga2*, and *YrU1* (Feuillet et al. 2003; Loutre et al. 2009; Wang et al. 2020a), stem rust resistance genes *Sr13* and *Sr8155B1* (Zhang et al. 2017; Shen et al. 2025), yellow rust resistance gene *TaRGA3* (Fang et al. 2024), and powdery mildew resistance gene *Pm41* (Li et al. 2020) (Fig. 2A). Among these, only Sr13 and its Kronos ortholog TrturKRN6A02G062130 shared identical protein sequences, while others showed minor sequence variations (Table S11).

**Figure 2.**
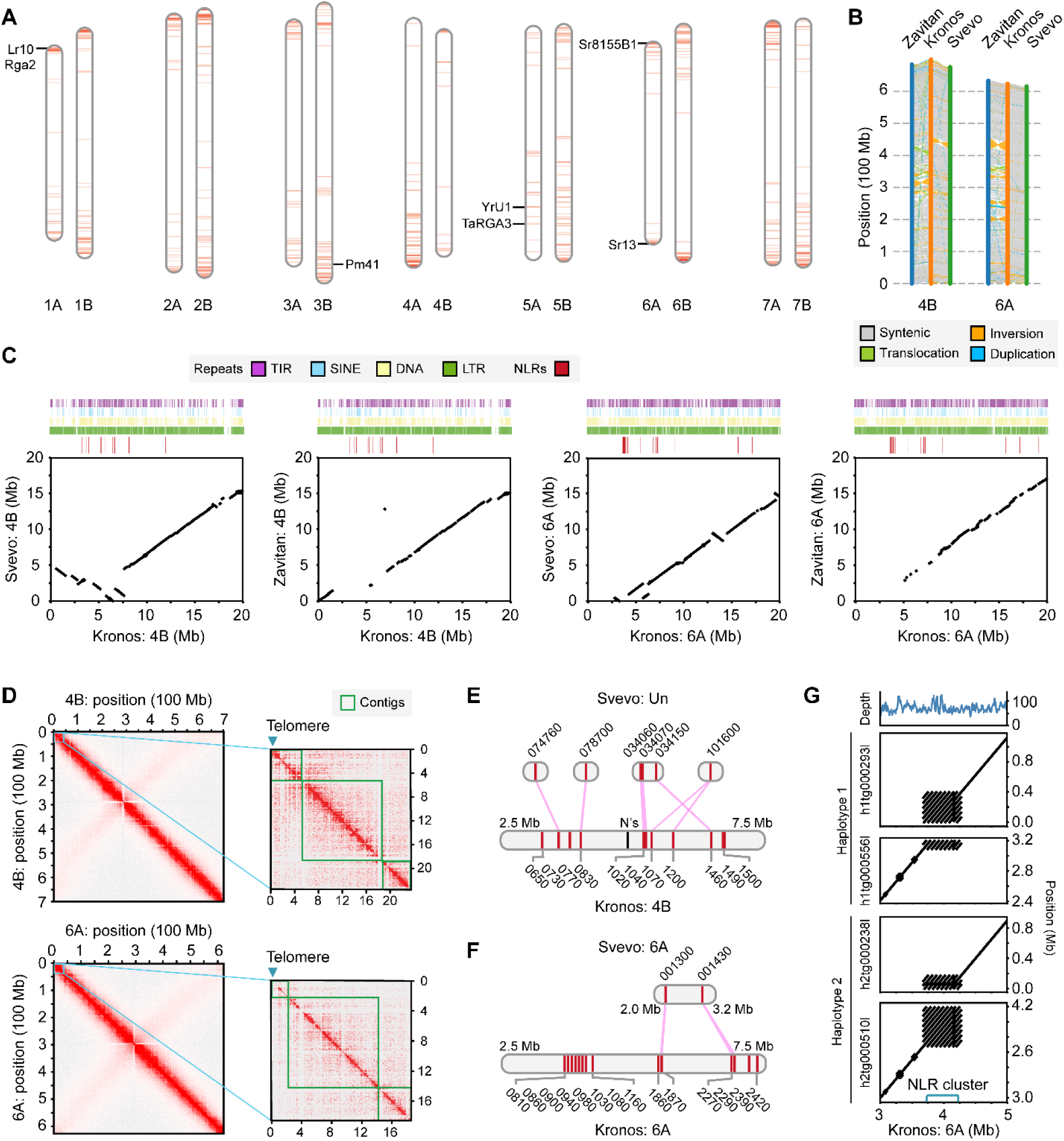
Genomic distribution, structural variation, and haplotype diversity of NLR clusters in the Kronos genome. **(A)** Chromosomal distribution of 1,089 reliable NLR genes across the Kronos genome, plotted in 3 Mb windows across the 14 Kronos chromosomes. Each ideogram is oriented from top (chromosome start) to bottom (chromosome end). Darker red indicates higher local NLR gene density. Positions of selected functional NLR orthologs are labeled. **(B)** Synteny of chromosomes 4B and 6A between Kronos, Svevo, and Zavitan. **(C)** Local synteny of the initial 20 Mb of chromosomes 4B and 6A between Kronos and Svevo or Zavitan. Tracks above show the four most abundant repeat types and reliable Kronos NLRs. Dot plots below show collinearity. **(D)** Hi-C contact maps for chromosomes 4B and 6A in Kronos. Heatmap intensity reflects chromatin contact frequency. Insets zoom into telomeric regions, with green boxes marking contig boundaries. **(E** and **F)** Genomic distribution of reliable Kronos NLRs (red lines) on the 2.5 to 7.5 Mb regions of chromosomes 4B and 6A, compared to putative orthologs identified in Svevo. Pink lines indicate orthology relationships. **(G)** Comparison between the Kronos reference genome and haplotype-resolved assemblies across the 3.0 to 5.0 Mb region of chromosome 6A. The top track shows HiFi read coverage. Dot plots display collinearity between reference and each haplotype.

### The Kronos genome improves the resolution of NLR genomic organization

To assess concordance with existing assemblies, we performed whole genome synteny analyses. Overall, Kronos showed high synteny with the available Svevo, *T. turgidum* Zavitan, and Chinese Spring genome assemblies (Figs. S9 and S10), with higher concordance to Svevo than Zavitan (Fig. 2B; Fig. S11). Svevo was assembled with short reads, making telomere-proximal regions challenging to resolve (Maccaferri et al. 2019). We examined whether the Kronos assembly improved the resolution of NLR-rich intervals by comparing the terminal 20 Mb of each chromosome with the corresponding regions in Svevo and Zavitan (Figs. S12-S15).

On chromosomes 4B and 6A, consistent collinearity with other assemblies began at positions 7.5 Mb and 5 Mb, respectively (Fig. 2C). By contrast, the NLR cluster intervals (3.3-6.7 Mb on 4B and 3.7-4.2 Mb on 6A) were fragmented or absent in Svevo and Zavitan. The Kronos genome is 88% repetitive and dominated by LTR retrotransposons (70%) (Table S12). High repeat contents persisted near chromosomal ends, with LTRs being the most abundant type (Fig. 2C). Notably, synteny breakpoints still coincided with NLR-rich blocks. To confirm genomic accuracy, we examined Hi-C contact maps (Fig. 2D; Fig. S3). Telomeric repeats occurred at the beginning of both chromosomes, and continuous contact diagonals traversed from the telomeric ends to the NLR clusters, supporting structural integrity. In addition, inter-contig Hi-C connectivity corroborated the orientation and placement of contigs.

We searched for putative orthologs of Kronos NLRs located between 2.5 and 7.5 Mb on chromosomes 4B and 6A (Fig. 2E). No orthologs were identified in Zavitan (RefSeq v2.1), consistent with the lack of synteny in these intervals (Fig. 2C). In Svevo, five orthologs with identical protein sequences were present on unanchored contigs from chromosome 4B, suggesting unresolved placement (Fig. 2E). On 6A, two orthologs were identified, including partially annotated TRITD6Av1G001300 (Fig. 2F). Collectively, these findings highlight that the Kronos assembly and curated NLR set sharpen chromosomal context and continuity of NLR-rich, telomere-proximal regions.

### Residual heterozygosity may remain in Kronos

We validated the NLR cluster at 3.7–4.2 Mb on chromosome 6A (Fig. 2F) by mapping HiFi reads to the reference. Several regions showed ∼2X sequencing depth (Fig. 2G), indicating unresolved haplotype-specific structural variation. To investigate, we generated haplotype-resolved assemblies (AABB) and compared them to the haplotype-collapsed reference. In the reference, the NLR cluster appeared in a single contig (Fig. 2F). By contrast, neither haplotype assemblies fully spanned the interval, and one haplotype showed putative duplications within the cluster (Fig. 2G). These findings indicated that the assembler did not completely resolve highly diverged, structurally complex NLR clusters between haplotypes that may represent previously described residual heterozygosity (Krasileva et al. 2017). The apparent continuity in the reference may reflect a simplified or partially collapsed representation of a single haplotype.

### Hidden diversity and domain innovation among Kronos NLRs

Canonical wheat NLRs comprise an N-terminal coiled-coil (CC), a central NB-ARC domain and a C-terminal leucin-rich repeat (LRR) domain (Baggs et al. 2017). A subset carries integrated domains (IDs), often homologous to host targets of pathogen effectors to bait them for immune activation (Kroj et al. 2016; Sarris et al. 2016). To investigate phylogenetic diversity and relationships among Kronos NLRs and cloned functional NLRs (Table S11), we constructed a phylogenetic tree (Fig. 3). Among 1,089 reliable NLRs, 89 (8.2%) contained PFAM-annotated IDs (Table S13). Consistent with prior work (Bailey et al. 2018), NLR-IDs were unevenly distributed across the phylogeny and appeared in three previously defined major integration clades (MICs 1, 2 and 3) (Fig. 3). MICs 2 and 3 primarily contained DDE superfamily endonuclease domains (PF13359) and BED zinc finger domains (PF02892), respectively, whereas MIC 1 had diverse IDs. We also observed additional integration clades (Fig. S16) with protein kinase (Pkinase; PF00069) domains, including Tsn1 (Faris et al. 2010), and with major sperm (MSP; PF00635) domains (Fig. 3). Across all Kronos NLRs, Pkinase was the most observed ID (34 genes), followed by DDE endonuclease (13 genes) and MSP (9 genes) (Table S14).

**Figure 3.**
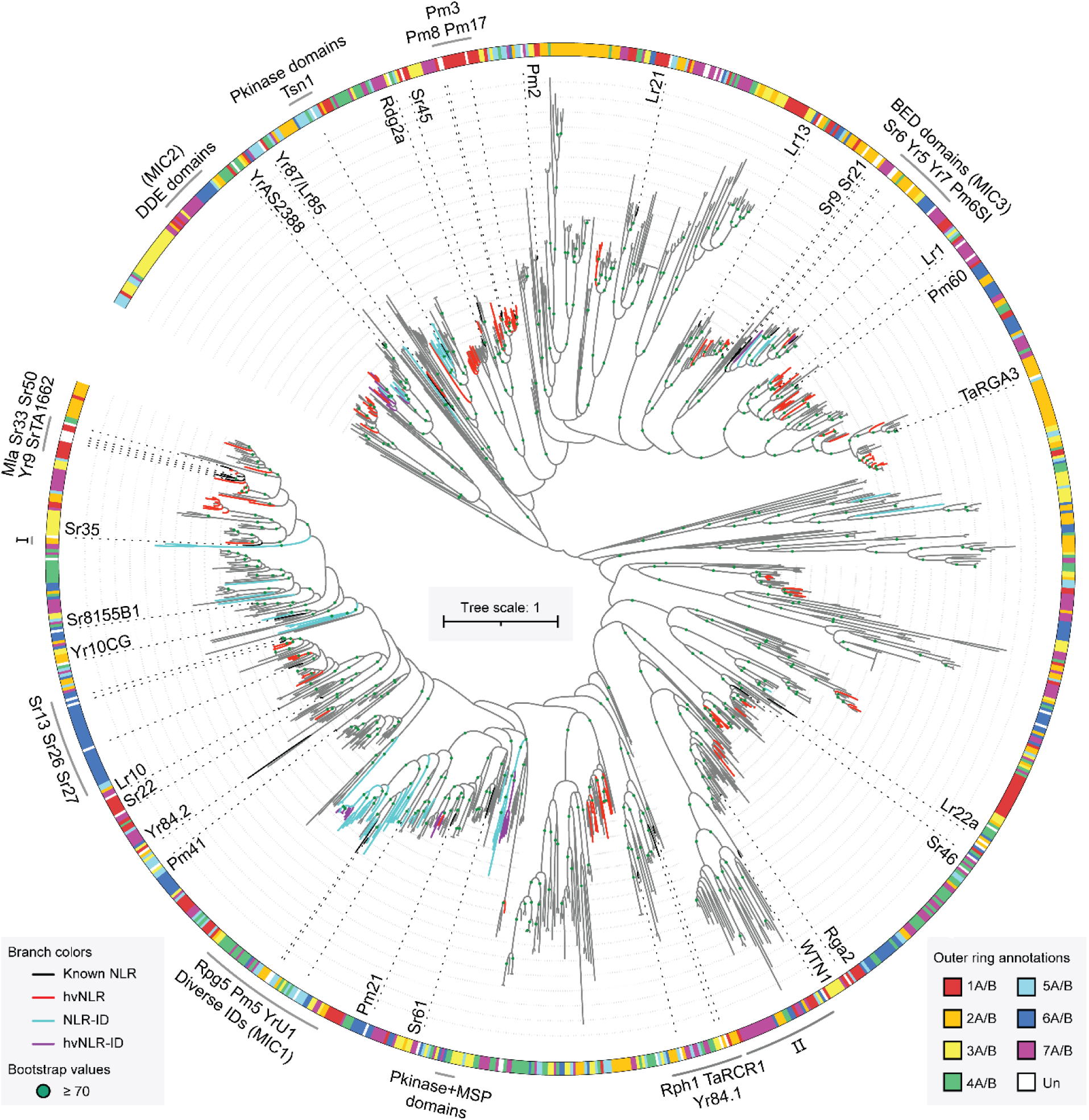
Phylogenetic tree of reliable Kronos NLRs and known functional NLRs. Maximum-likelihood phylogeny inferred from NB-ARC domains. Branch colors indicate: black—cloned, functionally validated NLRs; red—Kronos highly variable NLRs (hvNLRs); sky blue—Kronos NLR with PFAM-annotated integrated domains (NLR-ID); purple—Kronos hvNLR-IDs. All other Kronos NLRs are shown in gray. Names of cloned genes are labeled along dotted leader lines stemming from corresponding branches or around the perimeter. The outer ring denotes the chromosome of origin. Bootstrap supports ≥70 are shown as green circles at the corresponding nodes.

Some NLR-IDs appeared divergent compared to public databases. For example, TrturKRN2B02G099270 with a C-terminal Pkinase domain lacked close ortholog in NCBI, with its best match (XXBI60639.1) sharing only 70% sequence identity. This NLR formed a phylogenetically isolated micro-clade with TrturKRN2A02G083980 that shares 63% identity (Fig. 3: Clade Ⅰ; Fig. S17A). TrturKRN2B02G099270 lies near the distal end of chromosome 2B (Fig. 4A). Its nearest physical neighbor, TrturKRN2B02G099510, is phylogenetically distant, sharing only 25% sequence identity along the NB-ARC domain (Fig. 3: Clade ⅠⅠ; Fig. S17B). Furthermore, despite clear transcriptomic support for both expression and gene structure (Fig. 4B; Fig. S18), TrturKRN2B02G099510 also lacks close orthologs in NCBI, with its best match (XBI97030.1) showing 65% identity. Notably, TrturKRN2B02G099510 does not have PFAM-annotated IDs but carries simple repeat insertions within the LRR domain (Fig. 4B; Fig. S19). AlphaFold 3 modeling suggested that these sequences may form LRR-like structures, albeit with limited confidence (Fig. S20) (Abramson et al., 2024). Both genes sit within a cluster of 16 pseudogenized NLRs (Fig. 4A), suggesting that intense birth–death turnover and domain innovation can proceed in phylogenetically distant NLRs in a genomically isolated locus.

**Figure 4.**
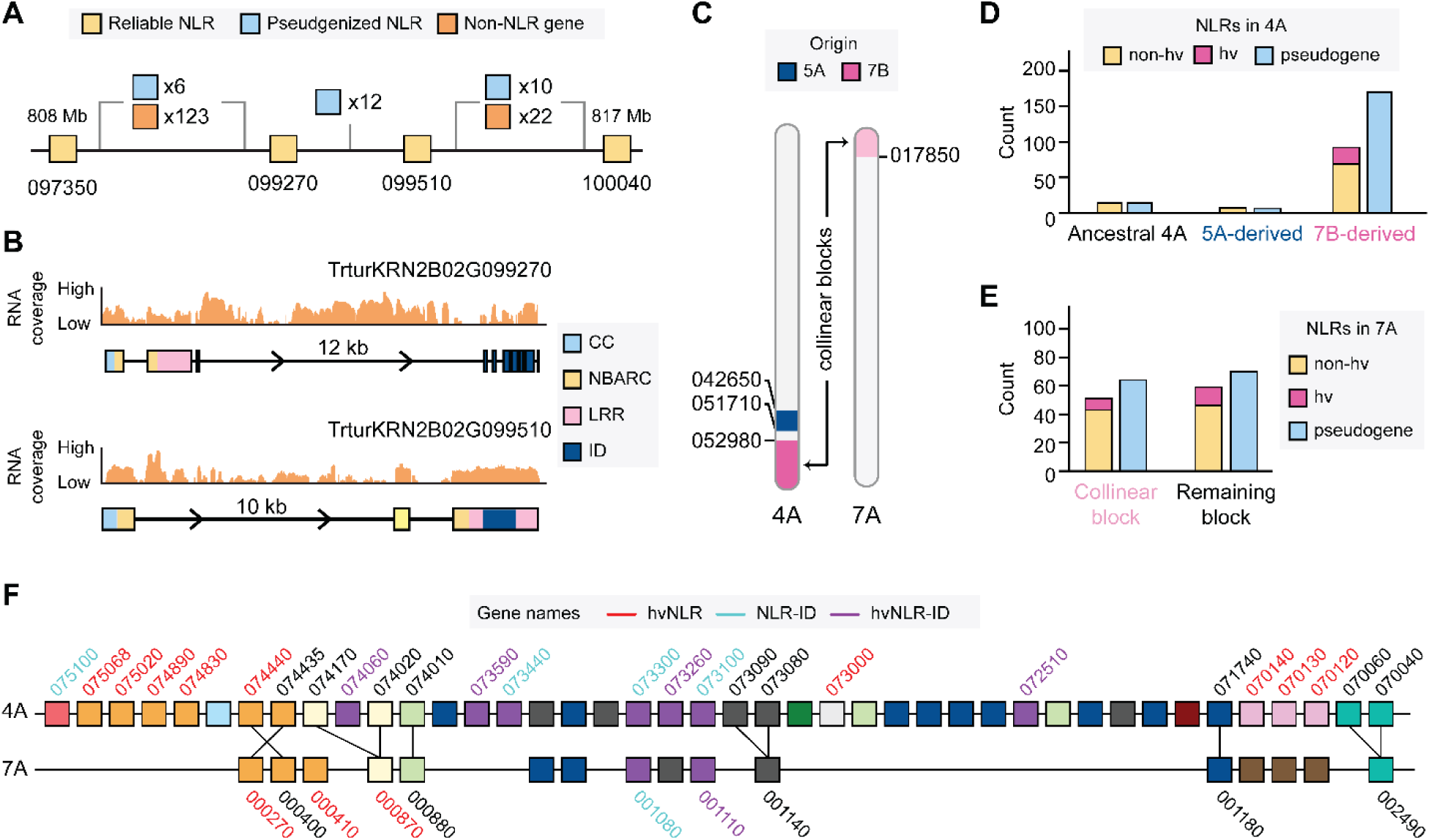
Structural and evolutionary diversity of Kronos NLRs. **(A)** Genomic organization of telomere-proximal NLR loci on chromosome 2B (808-817 Mb). Boxes indicate reliable NLRs (yellow), pseudogenized NLRs (sky blue), and non-NLR genes (orange). The number of genes is indicated between two reliable NLRs. **(B)** Transcriptomic evidence supports expression and annotation of two divergent NLRs within this cluster. RNA-seq coverage across TrturKRN2B02G099270 (top) and TrturKRN2B02G099510 (bottom) is shown. Domain architecture includes coiled-coil (CC; cyan), NB-ARC (yellow), leucine-rich repeat (LRR; green), and integrated domains (ID; purple). **(C)** The origin of chromosome 4A. Colored blocks denote chromosomal segments derived from 5A (blue) and 7B (pink). Gene numbers near breakpoints are indicated. Homologous regions on chromosome 7A identified by collinearity analysis are indicated in light pink. **(D** and **E)** Distribution of NLRs across chromosomal blocks. Numbers of reliable NLRs (hvNLRs in pink and non-hvNLRs in yellow) and pseudogenized NLRs are indicated in defined chromosomal blocks in 4A and 7B. **(F)** Organization of NLRs in chromosomes 4A and 7A. Reliable NLRs are shown as blocks colored by phylogenetic placement. Gene IDs are labeled for hvNLRs (red), NLR-IDs (sky blue), hvNLR-IDs (purple), and putative orthologs (linked by lines).

Apparent novelty in Kronos can also reflect annotation gaps in other genome assemblies rather than true absence. To contextualize these findings, we predicted ∼42,000 NLRs across 40 *Triticum* assemblies and searched for homologs of Kronos NLR-IDs (Table S15). The prediction of these genes partly relied on ab initio parameters trained on high-confidence Kronos NLRs that performed well in internal validation (Table S16); however, cross-species generalizability was not formally assessed due to limited manually curated test sets in other wheat species. For a substantial subset of Kronos NLR-IDs, including TrturKRN2B02G099270, we identified identical or near-identical orthologs (Table S17). Thus, the perceived uniqueness of these loci likely stems from mis- or under-annotation in prior references, with manual curation in Kronos revealing naturally occurring variants that were previously uncaptured.

### 7B-derived segment on chromosome 4A shows biased NLR accumulation and accelerated diversification

The contemporary chromosome 4A in wheat has a complex evolutionary history shaped by multiple chromosomal rearrangements, including a reciprocal translocation from chromosomes 5A and 7B that contributed to its long arm (Devos et al. 1995; Clavijo et al. 2017). Homologous chromosomes 4A and 4B differ sharply in their NLR content: chromosome 4B has the fewest NLRs in the genome, whereas chromosome 4A carries the greatest number of NB-ARC–encoding loci but also the highest proportion of NLR pseudogenes (63%) (Table S10). To trace the evolutionary origin of these loci, we assessed gene-based collinearity between the Kronos 4A and the D-genome progenitor *Aegilops tauschii*, resolving breakpoints for the 5A- and 7B-derived segments (Fig. 4C; Fig. S21) (Luo et al., 2017). This analysis placed TrturKRN4A02G052980 as the proximal boundary of the 7B-derived segment, closely matching a previously reported breakpoint (Table S18) (Dvorak et al. 2018). Within this 100 Mb tract, we annotated 262 NLR loci, including 170 (65%) pseudogenes or fragments, whereas the remaining 650 Mb of 4A contained only 41 loci (Fig. 4D).

As expected from evolutionary history, a collinearity block is present between chromosomes 4A and 7A (Fig. 4C; Table S19). This block on 7A contains 51 reliable NLRs, substantially fewer than the number encoded by the 7B-derived segment on 4A (Fig. 4E; Table S20). NLRs are known to evolve at variable rates, with a subset undergoing rapid diversification to maintain effector recognition specificity. These fast-evolving loci, termed highly variable NLRs (hvNLRs), were identified by quantifying interspecies sequence diversity using Shannon entropy and NLRs annotated from 40 *Triticum* genomes (Fig. 3; Table S21) (Prigozhin and Krasileva 2021). Of the 23 hvNLRs on 4A, all were confined to the 7B-derived segment (Fig. 4D). By contrast, only 8 of 21 hvNLRs on 7A were within the collinear block (Fig. 4E).

Phylogenetic analysis revealed that NLRs from the 7B-derived segment of 4A are dispersed across the tree but show localized clustering and co-occurrence with NLRs from 7A (Fig. S22). Yet, direct comparison of the two chromosomes highlighted contrasting evolutionary trajectories at their terminal region (Fig. 4F). Within this compared interval, 42 reliable NLRs spanning 12 phylogenetic clades were found on 4A, while 16 NLRs across 8 clades were present on 7A (Table S20). The 4A segment also contained a richer diversity of NLR-IDs and more hvNLRs. These expanded NLR-IDs (purple boxes; Fig. 4F) belonged to MIC 1 (Fig. 3), indicating lineage-specific proliferation. Together, these results demonstrate that the 7B-derived block experienced biased NLR accumulation and accelerated diversification after its translocation to chromosome 4A.

### Kronos reference enhances sensitivity and accuracy of EMS mutation calls

Kronos has extensive EC and PC re-sequencing datasets for EMS-mutagenized populations, but without a reference genome and high-quality annotation, full functional potential of these resources had remained incomplete. To improve resolution and accuracy, we remapped publicly available EC and PC data from 1,439 and 1,556 Kronos lines to the reference, respectively (Tables S22 and S23). High-confidence mutations were then identified using the MAPS pipeline and re-evaluated for their functional impacts, consistent with the approaches in prior studies (Krasileva et al. 2017; Zhang et al. 2023b).

Across EC data, the number of EMS-induced mutations identified in our study were broadly consistent with Krasileva et al. (2017), but with a modest increase in mutation counts (Fig. 5A; Table S24). Specifically, across 1,416 shared mutant lines analyzed under the high confidence threshold (HetMinCov5HomMinCov3), we detected an average of 2,832 mutations per line, a 5% increase over 2,697 previously reported (Table S24) (Krasileva et al. 2017). By contrast, non-EMS-type mutation calls were reduced by 23% (Fig. 5B; Table S24), indicating improved specificity afforded by the Kronos genome. While direct coordinate-based comparison is confounded by the fragmented nature of the early Chinese Spring reference genome (Krasileva et al. 2017), the higher contiguity and accuracy of the Kronos genome likely enabled recovering previously undetectable mutations in missing, poorly assembled or divergent regions.

**Figure 5.**
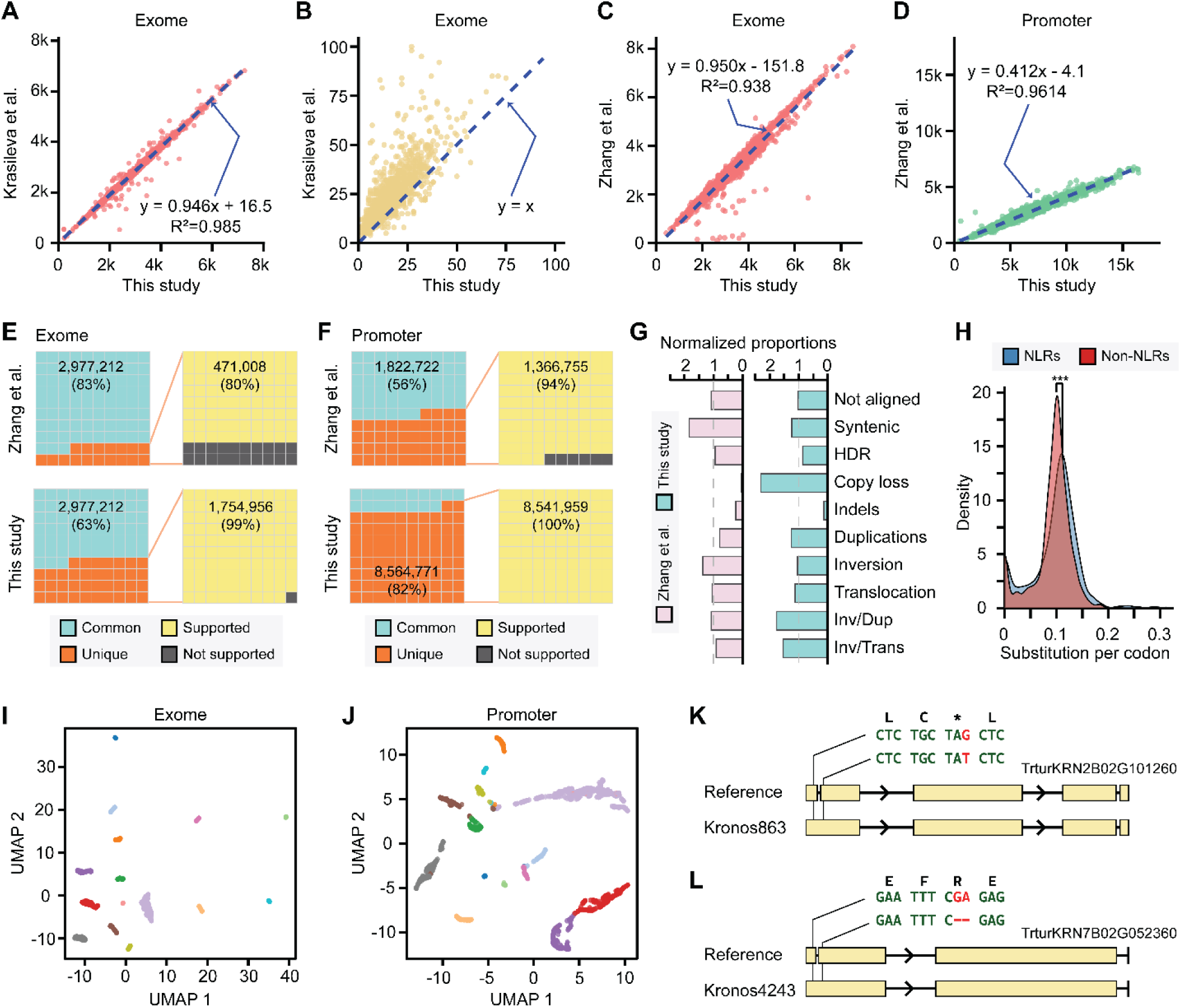
The Kronos reference genome expands the resolution of mutational landscapes of Kronos EMS mutants. **(A** and **B)** Comparison of EMS-type and non-EMS-type transitions, respectively, identified from exome-capture (EC) sequencing data of 1,416 Kronos mutants between this study and the previous study by Krasileva et al. (2017). **(C** and **D)** Comparison of EMS-type transitions identified from EC and promoter-capture (PC) sequencing data of 1,403 and 1,511 Kronos mutants, respectively, between this study and the previous study by Zhang et al. (2023). (**E** and **F**) Shared and unique mutations identified using the MAPS pipeline between this study and the study by Zhang et al. (2023). Unique mutations were further evaluated for support using GATK-based variant calling. **G.** Normalized proportions of mutation count by genomic category. Mutations uniquely identified from Zhang et al. (2023) and this study were assigned with genomic categories identified by SyRi. For each category, the EC-to-PC mutation count ratio was normalized by the total EC-to-PC mutation ratio, highlighting enrichment or depletion of each category. HDR: highly diverged regions; Inv: inversion; Dup: duplication; and Trans: translocation. **H**. Density distribution of high-confidence EMS-type substitutions per codon in NLRs and non-NLRs. These mutations were derived from both uniquely and multi-mapped reads. Non-NLR genes were randomly sampled to match the length distribution of NLRs. \*\*\**P* = 7 * 10^-6^ based on Wilcoxon rank sum test with continuity correction. **(I** and **J)** UMAP visualization of non-EMS-type mutations from EC and PC datasets, respectively. Mutations and zygosity states for non-EMS-type mutations present in at least 5% (EC) and 15% (PC) of mutants were included. The resulting mutation matrix was reduced using principal component analysis, and the top 50 principal components were used to construct UMAP embedding. Colors indicate cluster assignments derived from the EC dataset and are applied consistently across both panels. **(K** and **L)** Examples of non-EMS-type mutations in Kronos mutants that restore proper gene structures of low-confidence NLRs.

Zhang et al. (2023) leveraged the updated Chinese Spring reference genome (v1.0) and refined EMS mutation detection parameters to enhance sensitivity. To directly evaluate the impact of the Kronos genome, we compared mutation calls for commonly analyzed 1,403 EC and 1,512 PC datasets, using the EMS98% error method (Zhang et al. 2023b). Our analysis recovered an average of 3,391 EMS-type transitions for EC datasets, a 10% increase over 3,071 reported previously (Fig. 5C; Table S24). For PC datasets, we identified an average of 6,882 mutations per line, representing a 2.4-fold (143%) increase relative to 2,834 in the prior study (Fig. 5D; Table S25). These results demonstrate that mapping to the Kronos genome substantially improves the sensitivity and resolution of EMS mutation detection, yielding a more complete view of induced variation, especially in promoter regions.

### Regulatory region diversity drives variability in EMS mutation detection

To examine whether discrepancies in PC mutation calls between our study and Zhang et al. (2023) arose from technical artifacts of the MAPS pipeline, we independently called mutations with a standard GATK-based workflow. Unlike MAPS, which processes mutants in batches of 24 to 48 to specifically identify EMS-induced mutations, the GATK pipeline analyzes each line independently and recovers the full mutational landscape. Overall, 99.4% (EC) and 99.8% (PC) of high-confidence EMS-type transitions we identified using the MAPS pipeline were consistently recovered by the GATK analysis, supporting the reliability of our results (Table S26).

To further investigate differences from Zhang et al. (2023), we lifted their reported mutations from the Chinese Spring genome to the Kronos genome. Among the EMS-type transitions, 84% (EC) and 76% (PC) could be successfully lifted over to Kronos coordinates (Table S27). Of these, 83% (EC) and 56% (PC) were consistently present in our MAPS-based calls (Fig. 5E and 5F). Given the larger discrepancy in PC, we examined whether our GATK calls could capture additional overlaps. Allowing a ±2 base pair window to account for small insertions and deletions, GATK recovered 80% (EC) and 94% (PC) of the unique variants retrieved by Zhang et al. (2023) (Fig. 5E and 5F). These results indicate that most uniquely detected mutations in both studies are likely biologically supported. Variability in MAPS-based mutation calls instead stems from reference choice, alignment qualities and batch-based mutation calling.

To contextualize these findings, we associated uniquely identified mutations from the MAPS with genomic features. Notably, about half were located within non-syntenic regions (Fig. S23). Investigating the ratio of EC-to-PC mutations, we found that the use of the Kronos reference genome enabled recovery of more promoter-associated mutations, in regions shaped by multiple structural rearrangements, such as inversion/duplication and inversion/translocation (Fig. 5G; Table S28). This likely reflected both greater sequence complexity and higher divergence in promoter versus exonic regions, amplifying variation in mutation detection across reference genomes and batch-processing methods.

### The mutational landscapes across NLRs

EC and PC rely on pre-designed baits derived from existing genomic sequences. At the time of probe design, only transcriptomic data and a highly fragmented genome were available for Kronos (Krasileva et al. 2013). Therefore, certain genic regions, particularly NLRs, may have been underrepresented or entirely missed. We quantified the number of MAPS-derived EMS-type mutations across the coding regions of 1,089 reliable NLRs and compared them to an equivalent number of randomly sampled non-NLR Kronos genes with matched protein length distributions (Fig. 5H; Table S29; Fig. S24). The distributions differed significantly: non-NLRs displayed narrower, sharper profiles, indicating lower variance. By contrast, NLRs showed broader distributions, though not an extreme binomial-like pattern, suggesting that capture efficiency was largely maintained. The maximum density of high-confidence mutations, including uniquely and multi-mapped reads, was 0.3 per codon. On the contrary, 135 NLR showed < 0.03 (Table S29). These NLRs were distributed across phylogeny, and when they co-occurred, we also observed short branch lengths, indicating very recent duplications and inability to resolve alignment ambiguity (Fig. S25). In total, 924 out of 1089 reliable NLRs had premature stop codons across 1,286 mutants (Table S30). In addition, 221 NLRs in 283 mutants had splicing site mutations predicted to disrupt translation (Table S30). Together, these provide sufficient coverage to investigate NLR functions.

Kronos seeds used to generate the TILLING population came from a breeding program and contained residual genetic heterogeneity (Krasileva et al. 2017). Dimensionality reduction of non-EMS-type mutations using Uniform Manifold Approximation and Projection (UMAP) revealed distinct sub-populations of Kronos mutants (Fig. 5I), likely reflecting the underlying diversity of the seed stock. These sub-clusters persisted when using non-EMS-type mutations from promoter regions (Fig. 5J), or EMS-type transitions (Fig. S26), suggesting that substantial baseline diversity remains within the mutagenized lines. We hypothesized that some NLRs with recent disruptive mutations, such as premature stop codons (TrturKRN2B02G101260) and frameshift mutations (TrturKRN7B02G052360), may remain intact in some individuals in the EMS population. Indeed, we found rare mutant lines in which these loci lacked disruptive mutations and were associated with non-EMS-type mutations (Fig. 5K and 5L; Table S31). Together, these findings demonstrate that the Kronos reference genome enables not only accurate mapping of EMS-induced mutations but also resolution of standing variation, revealing the full mutational landscape across a genetically diverse population.

### A subset of NLRs is constitutively expressed across developmental stages

NLR regulation is typically best characterized under pathogen challenge, but available datasets for Kronos capture only developmental stages (Table S32). To establish a developmental baseline, we re-quantified all available RNA-seq datasets against the v2.1 annotation (Tables S33–43). Because direct comparison of counts across independent studies is confounded by batch effects, we transformed expression values into within-sample percentile ranks, assigning each gene a value from 0 (lowest expression) to 1 (highest expression) (Tables S44-46). Principal component analysis of NLR percentile ranks revealed clear tissue-specific clustering across 7 studies (Fig. 6A; Table S44). For instance, meristem datasets included wild type, *vernalization1* (*vrm1*), and *vrm1/fruitful2* (*ful2*) mutants and spanned both vegetative and developing shoot apical meristem (SAM) across three independent studies (Li et al. 2021; VanGessel et al. 2022; Xu et al. 2025). Nevertheless, these samples clustered together and were clearly separated from seed and leaf (Fig. 6A). At a finer scale, however, mutants with impaired meristem development (*vrn1* and *vrn1/ful2*) showed reduced Pearson correlations to wild type (Fig. S27). Together, these results indicated that tissue identity is the primary determinant of NLR expression patterns in the absence of pathogen induction, with genotype introducing secondary structure within meristem samples.

**Figure 6.**
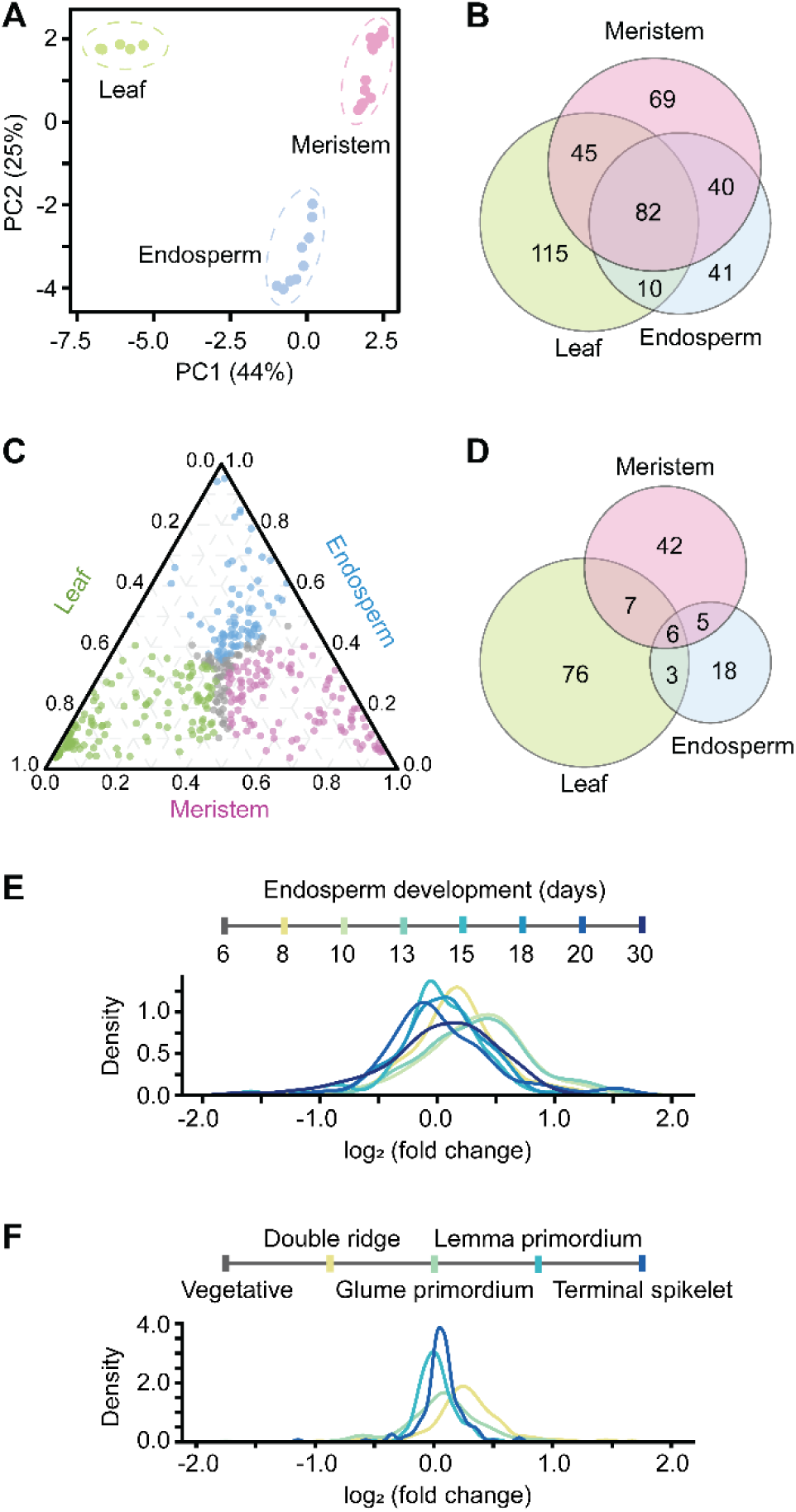
Constitutive expression of NLRs during wheat reproductive development. **(A)** Principal component analysis of transcriptome-wide percentile ranks for NLRs. Tissue types are indicated in colors: leaf (green), combined vegetative to terminal inflorescence meristem series (magenta), and endosperm (blue). **(B** and **D)** Venn diagrams displaying overlaps of NLRs with median percentile ranks ≥0.40 and ≥0.70, respectively. For meristem, only wild type samples were included for analysis. **(C)** Ternary plot of median global percentile rank distributions across tissues, with each point colored by the tissue of highest relative expression. **(E** and **F)** Density plots of log₂ (fold changes) across developmental stages. Colors indicate the focal stage being compared to the immediately preceding stage, illustrating relative shifts in expression from vegetative through terminal reproductive development.

To capture broad tissue-level trends, we summarized NLR expression as median percentiles per tissue and compared expressed sets at two thresholds: 0.40 and 0.70. At these cutoffs, normalized counts per million (CPM) corresponded to 1 to 7.5 and from 8.8 to 21.4, respectively, which displayed consistency within individual studies but variable across studies (Table S47). Across leaf, endosperm, and SAM, 402 NLRs were found at the 0.40 cutoff, with 82 shared across all tissues (Fig. 6B; Table S48). At the higher 0.70 threshold, a distinct and substantial set of 76 NLRs showed preferential expression in leaves, indicating their likely exposure to pathogen challenge in this tissue (Fig. 6C and 6D). Unexpectedly, 42 NLRs were identified only in SAM (Fig. 6D; Table S48), with meristem-specific high expression (Fig. 6C). Phylogenetic mapping of these NLRs revealed no notable clade-specific enrichment, indicating that baseline activity is broadly distributed rather than restricted to particular clades (Fig. S28).

Using stage-resolved RNA-seq datasets (VanGessel et al. 2022; Chen et al. 2023), we tracked the expression dynamics of 173 and 236 NLRs with median percentiles ≥0.40 in endosperm and SAM, respectively (Fig. 6B). In endosperm, most NLRs maintained stable expression across development (Fig. 6E): median relative expressions between successive stages were close to zero, and >90% of genes remained above -0.5 log fold changes, indicating that very few NLRs were transcriptionally silenced. A similar pattern was observed in SAM, but with tighter expression controls than endosperm (Fig. 6F), suggesting that reproductive development preserves a stable baseline of NLR activity by deploying distinct yet partially overlapping NLR sets. This persistence possibly suggested either continuous immune surveillance as an intrinsic feature of wheat development or potentially new and yet to be uncovered roles of NLRs in SAM.

### Annotation of miRNA and phasiRNA loci in Kronos establishes a framework for sRNA-based studies

To support developmental studies, we curated *MIR* and *PHAS* loci in the Kronos genome by reanalyzing sRNA libraries collected from anthers (Bélanger et al. 2025) (Table S49). All 157 miRNA loci previously identified from Svevo were successfully transferred to Kronos, including 134 loci from 26 conserved miRNA families and 23 candidate loci (Fig. S29; Table S50). Only 9 loci showed minor sequence variations (98.5% to 99.9%), highlighting strong conservation of miRNA-producing loci between the two genomes. Consistently, *MIR* order was collinear across the two genomes (Fig. S29), except for a short interval on chromosome 3A characterized by a local inversion (Fig. S30).

In Kronos, we identified 7,662 21-*PHAS* and 3,303 24-*PHAS* loci mostly distributed along chromosomal ends, compared to 6,405 and 2,699 loci in Svevo (Fig. 7A) (Bélanger et al. 2025). Both 21-nt and 24-nt phasiRNAs displayed developmental accumulation in anthers (Fig. S31A). By contrast, 24-nt phasiRNAs were completely depleted across all stages of anther development in male-sterile *dicer-like 5 (dcl5)* mutants (Fig. S31B). Among putative *PHAS* precursors, 90.8% of 21-*PHAS* contained the conserved miR2118 trigger motif (Table S51). Known motifs were also present in 24-*PHAS* precursors, with the AU-rich motif in 81.9% of premeiotic precursors and the miR2275 motif in 87.3% of meiotic precursors (Table S51). Genomic analysis showed almost no overlaps between *PHAS* loci and protein-coding genes, only with 55 and 37 overlaps with 21-*PHAS* and 24-*PHAS* loci. This indicated that almost all *PHAS* loci originate from non-coding regions transcribed by RNA Polymerase II. Collectively, these results provide high-confidence annotation of *MIR* and *PHAS* loci in the Kronos genome and establish a robust framework for sRNA-based studies in Kronos.

**Figure 7.**
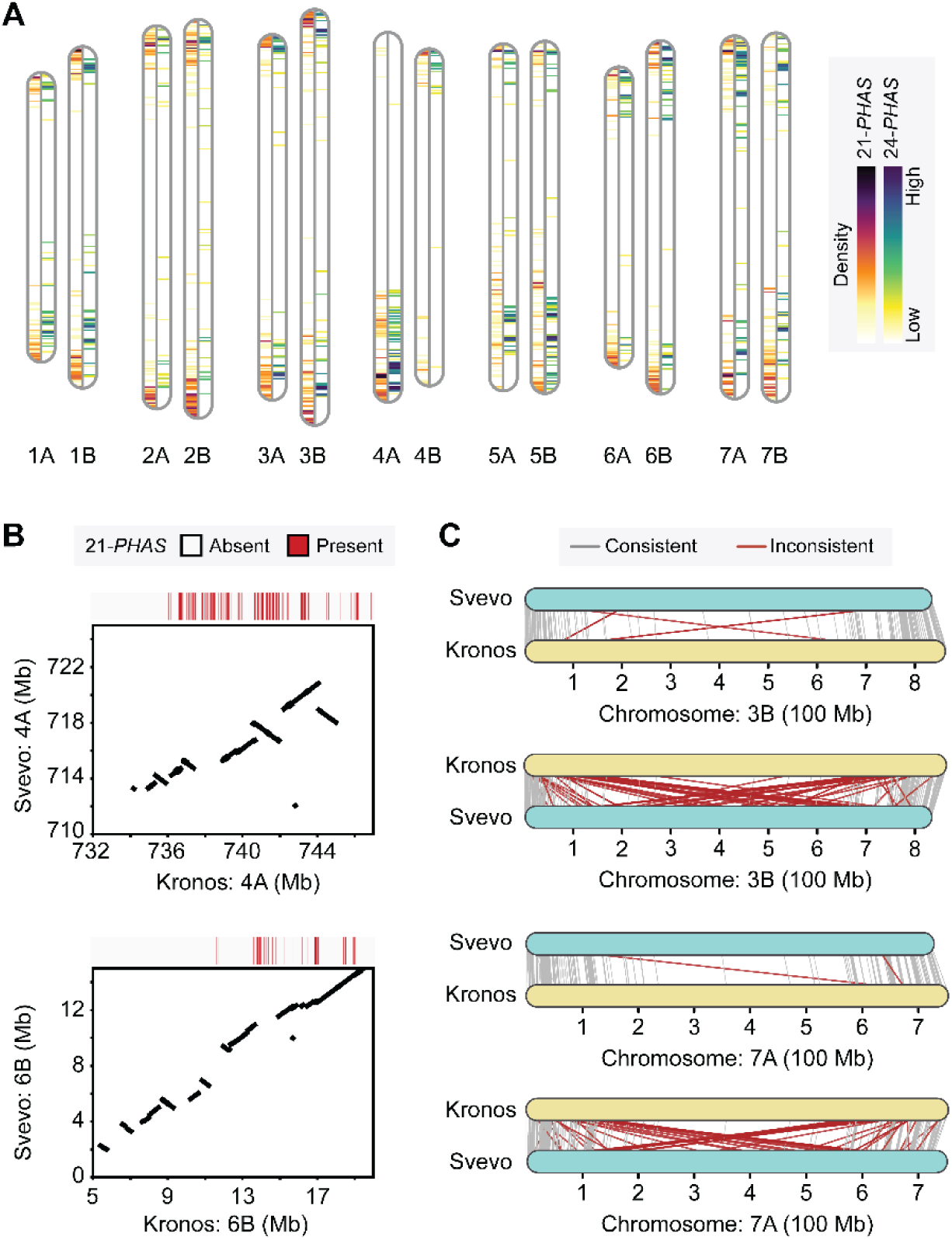
Newly annotated *PHAS* loci are associated with Kronos-specific genomic variations. **(A)** Distribution of 7,662 21-*PHAS* and 3,303 24-*PHAS* loci across the Kronos genome, shown using a 3 Mb sliding window. Each ideogram represents a chromosome, oriented from top (chromosome start) to bottom (chromosome end). **(B)** Local synteny of the selected 15Mb regions in chromosomes 4A and 6B between Kronos and Svevo. The top tracks indicate the presence of 21-*PHAS* loci in the Kronos genome. Dot plots show collinearity. **(C)** *PHAS* loci annotated in one genome (top) were mapped to the corresponding chromosome in the other genome (bottom). Each line connects homologous positions: grey lines indicate consistent liftover between genomes, while red lines indicate inconsistent positioning. Two chromosomes, 3B and 7A, are selected as examples.

### Unique *PHAS* loci associate with genomic variations

The Kronos genome has more annotated *PHAS* loci than the currently available Svevo genome. Comparative analysis revealed that many of the additional loci in Kronos were in genomic regions associated with breaks in synteny (Fig. 7B; Table S52). Reciprocal similarity searches confirmed that nearly all *PHAS* loci annotated in Svevo were also present in Kronos: 9,104 Svevo-derived precursors produced 9,104 top placements in Kronos, with 9,080 (99.7%) mapping to syntenic chromosomes, consistent with strong macro-synteny between assemblies (Fig. 7C; Fig. S32). In contrast, the reciprocal search from Kronos to Svevo yielded a higher rate of inconsistent placement (Fig. 7C; Fig. S33), possibly indicating their association with non-syntenic or Kronos-specific sequences. Together, these results demonstrate that genome-specific structural variation underlies part of the expanded *PHAS* annotation in Kronos, underscoring the importance of high-quality references for comprehensive annotation of sRNA loci.

## Discussion

We present a chromosome-scale reference genome for *T. turgidum* cv. Kronos with near-complete chromosomal continuity, high quality annotations for both protein-coding genes and sRNAs, and integrated functional genomic resources. A key advance is the resolution of telomere-proximal regions, where other currently available *T. turgidum* genomes show limitations. This contiguity allows accurate placement of NLR-rich intervals and finer analyses of their organization, expression, copy number, and diversity. Reanalysis of community resources, particularly TILLING exome capture and promoter capture datasets, uncovered mutations previously hidden in exonic and promoter regions, including 2.4-fold increase in promoter mutation calls. Similarly, re-annotation of *MIR* and *PHAS* loci yielded a more comprehensive, chromosome-wide catalog of sRNAs. Together, these advances position Kronos as a model plant for basic and applied research.

NLRs not only underpin disease resistance in crops but also exemplify the dynamics of rapidly diversifying genomic loci and offer templates for engineering new recognition specificities. Our manually curated NLR repertoire highlights hidden diversity and offers an anchoring framework for discovering functional immune receptors and annotating NLRs in related wheat species. High-quality long-read assembly and this annotation were essential to reveal fast evolving regions, such as the 7B-derived segment on chromosome 4A, where NLRs displayed accelerated diversity regeneration, by both point mutations and structural variations. These insights into NLR evolution can guide both comparative studies and targeted cloning of new functional alleles. Kronos had intermediate adult resistance to previously predominant races of stripe rust at the adult plant stage. At the seedling stage, it is fully susceptible to stripe rust, the Septoria tritici blotch, and leaf rust diseases. However, Kronos has seedling resistance to tan spot and intermediate seedling resistance to powdery mildew in greenhouse tests, providing a source of yet-uncloned resistance genes (Lunde et al. 2025). Their identification is now enabled by our genomic resources.

Notably, we observed that many NLRs were expressed across vegetative and reproductive tissues, including SAM. Previous work has shown that NLRs are stably expressed in roots and shoots in expectation of pathogen challenges, with monocots displaying preferential expression in roots (Munch et al. 2018). Recent studies have further demonstrated that NLRs, including well characterized functional receptors, are constitutively expressed in the absence of pathogens (Brabham et al. 2024; Prigozhin et al. 2024; Sutherland et al. 2024, 2025), and high expression in leaves can be used to prioritize candidates for functional testing (Brabham et al. 2024). In contrast, reproductive tissues generally display depleted NLR expression (Munch et al. 2018), consistent with the hypothesis that NLR activity is suppressed to minimize the risk of autoimmunity. Importantly, constitutive transcript accumulation does not necessarily equate to functional surveillance, since NLR activity is subject to post-transcriptional regulation. Elevated expressions may instead increase the mutation likelihood in anticipation of future pathogen encounters (Staunton et al. 2023). Alternatively, the presence of distinct NLR sets in SAM may point to uncharacterized functions. The meristem is remarkably resistant to pathogen invasion, with immune pathways that restrict pathogen proliferation while preserving stem cell integrity (Wu et al. 2020a, 2020b). Our observation raises the intriguing possibility that these receptors contribute to the immune resilience of this developmental niche.

In line with the collaborative history of wheat genomics, we have prioritized open distribution of resources. Beginning in November 2023, all primary and derived datasets were released on Zenodo with versioned DOIs, nearly two years before the completion of this study (Table S1). Since their release, the Kronos genome and associated functional resources have been downloaded over 2,500 times, prompting community requests and accelerating discovery in wheat genomics (Dang et al. 2025; Xu et al. 2025). We expect this integrated functional genomics resource to continue supporting the development and curation of future wheat genomes (Peters Haugrud et al. 2025) and the use of wheat as a model system across research communities.

## Materials and Methods

### PacBio sequencing

High molecular weight (HMW) Kronos (PI576168) genomic DNA was extracted from 5 g of juvenile leaf and immature inflorescence tissue from a single plant. We used the Nanobind plant nuclei big DNA kit, following manufacturer instructions for grinding in liquid nitrogen (102-574-800). Femto Pulse was performed using a Genomic DNA 165 kb Kit (Agilent Technologies) for unsheared gDNA to ensure the quality of HMW DNA sizes. Two HiFi long-read sequencing libraries were prepared from 30 µg of gDNA. These libraries were sequenced on 6 PacBio REVIO 25M SMRT cells by the DNA Technologies and Expression Analysis Core at the University of California, Davis. The resulting bam files containing HiFi reads were converted to the fastq format with bam2fastq v3.0.0, a module in the pbtk package (https://github.com/PacificBiosciences/pbtk).

### Chromosome conformation capture sequencing

Three proximity ligation libraries were prepared, with each using 500 mg of juvenile leaf and immature inflorescence tissue. These materials came from a single sibling of the plant used for the PacBio library preparation. Library construction followed the manufacturer’s protocol (Proximo Hi-C Plant Kit KT3045, version 4.5; Phase Genomics, Inc). DpnII (GATC), DdeI (CTNAG), HinfI (GANTC), and MseI (TTAA) restriction enzymes were used for digestion in all three libraries. Each library was sequenced as 150 bp paired-end reads in NovaSeq 6000 through Phase Genomics. Adapter and poly-G sequences were trimmed, and any reads with a quality score less than 20 or a length less than 50 were discarded by fastp v0.23.2 (-g --detect_adapter_for_pe -q 20 -l 50) (Chen et al. 2018). These filtered reads were used for scaffolding.

### Genome assessment

We assessed the genomic characteristics of Kronos using GenomeScope v2.0 (Ranallo-Benavidez et al. 2020). K-mers (21-mers) were analyzed from HiFi reads with Jellyfish v2.2.10 (-C -m 21), and a histogram was generated (-h 5000000) (Marçais and Kingsford 2011). GenomeScope was run on the histogram for tetraploidy (-p 4). For comparison to the Svevo genome, we downloaded paired-end Illumina sequencing data from PRJEB22687, trimmed the reads with trim_galore v0.6.6 and cutadapt v3.7 (--illumina) (Martin 2011), and repeated the GenomeScope analysis on the filtered reads.

### Genome assembly and scaffolding

We used hifiasm v0.19.5-r587 to assemble a haplotype-collapsed genome (-l0) or haplotype-resolved genomes with integration of Hi-C datasets (Cheng et al. 2021). For the collapsed genome assembly, a small size of associate contigs were concatenated with primary contigs to create an initial assembly. We followed the Omni-C pipeline for scaffolding (https://omni-c.readthedocs.io). The filtered paired-end Hi-C reads were mapped to the initial assembly with bwa v0.7.17-r1188 (-5SP -T0) (Li 2013). With pairtools v1.0.2 (Open2C et al. 2024), ligation pairs were searched from the alignments (--min-mapq 40 --walks-policy 5unique --max-inter-align-gap 30), and PCR and optical duplicates were removed. The filtered alignments were sorted with samtools v1.15.1 (Li et al. 2009) and processed with yahs v1.2a.2 to scaffold the initial assembly (-e GATC, GANTC, CTNAG, TTAA) (Zhou et al. 2023). The Hi-C contact map was generated with juicer v1.9.9 and visualized with Juicebox v2.20.00 (Durand et al. 2016).

The final scaffolds were compared to the Chinese Spring reference genome as well as chloroplast (NC_002762.1) and mitochondrial (NC_036024.1) genomes with minimap v2.24-r1122 (-x asm5) (IWGSC et al. 2014; Li 2018). Plasmids sequences were separated, and the 14 largest scaffolds were renamed, following the notation of the reference wheat genome. A small number of unassigned contigs were concatenated with 300 N’s and placed together into a single scaffold.

### Synteny analyses

For global synteny comparison, genomes were compared using minimap v2.24-r1122 and analyzed with SyRI v1.7.0 and plotsr v1.1.0 (Li 2018; Goel et al. 2019; Goel and Schneeberger 2022). Gene-based collinearity analyses were performed using all-vs-all protein similarity searches of the longest isoforms per genes with diamond v2.1.9 and MCScanX v1.0.0 and visualized with JCVI v1.5.7 (Wang et al. 2012; Buchfink et al. 2021; Tang et al. 2024). To analyze local genomic variants, pairwise genome comparisons were performed using BLAST v2.15.0 (Camacho et al. 2009).

### Repeat annotation

Repetitive elements were initially annotated using HiTE v3.0.0, and this repeat library was used by RepeatMasker v4.1.5 to soft-mask the reference genomes v1.0 and v1.1 (Smit et al. 2013; Hu et al. 2024). After generating the reference annotation v2.0, we re-annotated repetitive elements with EDTA v2.2.2 (Ou et al. 2019). To enhance repeat prediction and classification, complete and consensus repeats for *Triticum* were retrieved from the TREP database and included as curated libraries (Schlagenhauf and Wicker 2016). Additionally, classified repeats from HiTE were integrated as RepeatModeler libraries. To prevent over-masking, the coding sequences of the v2.0 annotations were also provided.

### Initial annotations for protein-coding genes (v1.0)

Annotation v1.0 was generated by integrating short-read paired-end sequencing datasets publicly available for Kronos with multiple gene prediction tools to produce consensus gene models. RNA-seq data were retrieved from the NCBI using sratoolkit v3.1.1 (https://github.com/ncbi/sra-tools) and trimmed with trim_galore v0.6.6 and cutadapt v3.7 (Martin 2011). Transcript assembly was performed using both genome-guided and de novo strategies. For genome-guided assembly, the filtered reads were mapped to the Kronos genome using hisat v2.2.1 (Kim et al. 2019), and primary alignments were selected using samtools v1.20, and transcripts were assembled with stringtie v2.1.7 (Li et al. 2009; Pertea et al. 2015). In parallel, de novo and genome-guided assembly was carried out with Trinity v2.15.1 (Haas et al. 2013). Transcript assemblies from Trinity and StringTie were integrated using PASA v2.5.3 to support gene model refinement (Haas 2003).

Initial gene models were predicted using three different software: BRAKER v3.0.6, Funannotate v1.8.15 and GINGER v1.0.1 (Palmer and Stajich 2023; Taniguchi et al. 2023; Gabriel et al. 2024). BRAKER incorporated transcriptome evidence and the Poales protein sequences from the UniProt database (The UniProt Consortium 2019). Funannotate used pre-trained AUGUSTUS and SNAP parameters and incorporated transcriptome evidence (Korf 2004; Hoff and Stanke 2019). Similarly, GINGER relied on pre-trained ab initio parameters generated with different training sets, transcriptomic evidence and protein evidence, including sequences from selected *Triticum* species obtained from Ensembl Plant (Bolser et al. 2015). All predicted gene models, along with PASA transcripts and protein evidence generated using Miniprot v0.12 by aligning 2.8 million Poales protein sequences to the genome (Li 2023), were integrated using EvidenceModeler v2.1.0 to produce consensus gene models (Haas et al. 2008). PASA was then used to add untranslated regions and alternative isoforms.

### Improved annotations for protein-coding genes (v2.0)

Annotation v2.0 incorporated publicly available long-read transcriptomic data for *Triticum* species. Datasets were downloaded from the NCBI using sratoolkit v3.1.1 and aligned to the genome with minimap v2.28-r1209 (Li 2018). Transcript assemblies were generated using stringtie v2.1.7 and Isoquant v3.5.2 (Shumate et al. 2022; Prjibelski et al. 2023). To enhance splice junction detection, both long-read assemblies were supplemented with the short-read data previously used in annotation v1.0. Assembled transcripts were evaluated and selected using Mikado v2.3.4 (Venturini et al. 2018). Extensive manual examination followed, involving visual inspection and cross-comparison across v1.0 gene models, mikado gene models, and annotations from BRAKER, GINGER, and Funannotate. This process focused on resolving overlapping gene models, mis-annotated UTRs, paralog fusions, and chimeric gene structures. Through numerous rounds of refinement, involving discarding, rescuing, and re-evaluating models, a curated set of 71,357 genes was selected for inclusion in annotation v2.0.

### Annotation completeness assessment

The completeness of the genome annotations was assessed using BUSCO v5.4.2 and the poales_odb10 database, as well as OMArk v0.3.0 and the LUCA.h5 database (Simão et al. 2015; Nevers et al. 2025).

### Functional annotations

Functional annotations were performed using eggNOG-mapper v2.1.12 and InterProScan v5.68.100 (Jones et al. 2014; Cantalapiedra et al. 2021).

### NLR manual curation

To facilitate targeted NLR curation, we first followed a modified version of the previous workflow (Seong et al. 2020). The Kronos genome was translated in six frames, and open reading frames (ORFs) were identified using orfipy v0.0.4 (Singh and Wurtele 2021). The predicted ORFs were searched for NB-ARC domains using the Hidden Markov Model (HMM) obtained from PFAM (PF00931) with hmmsearch v3.4 (--domE 1e-4 -E 1e-4) (Eddy 2011; Mistry et al. 2021). While this approach captured the majority of putative NLR loci, some divergent NB-ARC domains remained undetected. To recover additional loci, we also applied NLR-Annotator v2.1b (Steuernagel et al. 2020). All loci identified by either method were extracted with 15,000 flanking sequences on both sides.

Initial gene models were predicted using MAKER v3.01.03 (Cantarel et al. 2008). Protein evidence included NLR sequences from 18 Poaceae species (Toghani and Kamoun 2024) and 415 reference NLRs from RefPlantNLR (Kourelis et al. 2021). Transcript evidence was provided by Stringtie assemblies of short-read and long-read RNA-seq data produced during v1.0 and v2.0 annotations. In addition, ab initio prediction models trained during v1.0 annotation were used, including Augustus parameters from BRAKER and SNAP parameters from GINGER.

Additional evidence was generated to enhance accuracy of curation. All intermediate and final gene models from reference genome annotations were incorporated. To assist with splicing junction annotations, selected RNA-seq data were mapped to putative NLR loci using STAR v2.7.11b (Dobin et al. 2013), and coverage profiles were generated using bamCoverage from deepTools v3.5.5 (Ramírez et al. 2014). Genomic sequences were translated into six frames, and domains were predicted with InterProScan v5.68-100.0 (Jones et al. 2014).

Predicted gene models and all evidence were loaded into Apollo Genome Browser v2.0.6 (Dunn et al. 2019). In the first round of curation, models containing NB-ARC domains were systematically evaluated through visual inspection of gene structures and domain architectures. Each predicted model was additionally queried against the NCBI non-redundant database to assess sequence homology and integrity of gene structures. In a second round of curation, gene models were re-examined and assigned confidence levels. Genes with splice sites supported by transcriptome data were designated high confidence. Genes lacking expression support were evaluated based on structural conservation with homologs supported by transcript evidence in NCBI. Genes with disrupted splice sites, frameshifts, or premature stop codons were classified as low-confidence, whereas models lacking strong evidence but without contradictory features were designated medium-confidence.

### Updated annotations for protein-coding genes (v2.1)

High- and medium-confidence NLR loci were incorporated into the v2.1 annotation set, while genes located in low-confidence loci were excluded.

### Phylogenetic analysis

A total of 1,089 reliable Kronos NLRs and 47 reference NLRs were aligned to the HMM profile of the NB-ARC domain with hmmalign v3.4 to capture homologous domain regions (Eddy 2011). The resulting sequences were re-aligned with MAFFT v7.525 (--globalpair –maxiterate 1000) (Katoh and Standley 2013). Gappy columns were removed with trimAl v1.5.rev0 (-gt 0.3) (Capella-Gutiérrez et al. 2009), and 15 sequences with >40% gapped sites were discarded. The final dataset contained 1,074 Kronos-derived NB-ARC domain sequences. A phylogenetic tree was constructed using RAxML-NG v1.2.2 (Kozlov et al. 2019) under the LG+G8+F substitution model. Tree inference included 50 independent tree searches from parsimony-based starting trees and 1,000 bootstrap replicates to assess node support. The phylogenetic tree was visualized in ITOL (Letunic and Bork 2021).

### NLR prediction across wheat genomes

High-confidence Kronos NLR gene models were used to train Augustus v3.4.0 and SNAP v2006-07-28 (Korf 2004; Hoff and Stanke 2019). For each of 40 *Triticum* genomes (Table S15), candidate NLR-encoding regions were identified by searching for NB-ARC domains in six-frame translated genomic sequences and by NLR-Annotator (Eddy 2011; Steuernagel et al. 2020). Candidate loci were repeat-masked with EDTA-derived libraries using RepeatMasker v4.1.2 (Smit et al. 2013), and ab initio gene models were generated using both Augustus and SNAP.

For evidence-based annotations, Iso-Seq transcriptomic data available from *Triticum* species were retrieved from NCBI, and sequences that contain NB-ARC domains were extracted. High-confidence NLR transcripts were defined as sequences displaying ≥90% coverage to reliable Kronos NLRs. For each genome, MAKER v3.01.03 was run twice (Cantarel et al. 2008). The first run incorporated high-confidence Iso-Seq-derived NLR sequences with the trained Augustus parameter. The second run used NB-ARC domain-containing sequences in the Iso-Seq datasets, the trained Augustus and SNAP parameters, and EDTA-derived repeat libraries. In both runs, reliable Kronos NLRs and curated functional NLR sequences from RefPlantNLR were used as additional evidence (Kourelis et al. 2021).

Gene models derived from four independent prediction strategies were compared to reliable Kronos NLRs by pairwise sequence alignments constructed with Diamond v2.1.9 (Buchfink et al. 2021). Alignments were inspected to identify pseudogene signatures such as disrupted splicing sites, or missing start and stop codons. Putative pseudogenes were excluded, and one representative model was retained per locus to minimize noise in downstream evolutionary analyses.

### HvNLR identification

We developed a fully automated Snakemake-based pipeline for identifying hvNLRs (Mölder et al. 2021). This workflow takes a collection of NLR sequences, constructs alignments and infers phylogenetic trees of NLR clusters. Phylogeny-based initial clustering in an earlier version was (Prigozhin et al., 2024) replaced by MMseqs2 with coverage mode of 1, identity threshold of 0.4 and coverage of 0.7 (Steinegger and Söding 2018). These parameters were chosen experimentally to preserve hvNLR clades in the published manually curated datasets (Prigozhin and Krasileva 2021; Prigozhin et al., 2024). Clustered sequences were aligned with MAFFT v7.520, and phylogenetic trees were built for each cluster using RAxML-NG v1.2.2 with model JTT and 100 bootstrap replicates (Katoh and Standley 2013; Kozlov et al. 2019). Each cluster was automatically refined in iterations. First, candidate tree branches were identified based on ecotype overlaps (minimum 10 ecotypes), maximum allowed branch length (0.3), and minimum required bootstrap support (90). Then, the tree was split in two by cutting the branch with the highest ecotype overlap. In case of multiple candidate branches, the longest branch was selected. Next, ecotype overlap was updated for the resulting sub-trees. This process was repeated until no further candidate branches remained. Lastly, alignment entropy scores for the sub-cluster alignments were calculated, and hvNLRs were defined as sequence clusters with ≥10 positions displaying entropy ≥2 bits. This workflow was used to analyze NLRs from the Kronos genome and 40 *Triticum* genomes.

### Non-coding RNA prediction

To identify lncRNAs, we processed two sets of transcripts: those assembled by stringtie in the v1.0 annotation and those derived from mikado labeled as ncRNAs in the v2.0 annotation. Transcripts shorter than 200 nucleotides or with a single exon were excluded. The remaining candidates were compared against the v2.1 annotation, and any models overlapping with annotated coding genes were removed. We followed the lncRNA identification workflow, Plant-LncPipe-V2 (Tian et al. 2024, 2025), which integrates predictions from PlantLncBoost, LncFinder, and CPAT, along with sequence similarity-based filtering using UniProt database, to distinguish non-coding from coding transcripts. Lastly, FEELnc v0.2 was used to classify lncRNAs with respect to adjacent coding genes in the v2.1 reference annotation (Wucher et al. 2017).

Small nucleolar RNAs (snoRNAs) and small nuclear RNA (snRNAs) were identified by searching against the Rfam v15.0 database using cmscan from Infernal v1.1.5 (Nawrocki et al. 2009; Ontiveros-Palacios et al. 2025). Transfer RNAs (tRNAs) were predicted using tRNAscan-SE v2.0.12 (Chan et al. 2021), and ribosomal RNAs (rRNAs) were annotated using barrnap v0.9 and hmmer v2.3.2 (https://github.com/tseemann/barrnap).

### *MIR* and *PHAS* loci analyses

We analyzed sRNA-seq datasets from premeiotic, meiotic and postmeiotic anthers of wild-type Kronos and *dcl5* mutant lines from PRJNA1204935 (Bélanger et al. 2025). Adapter trimming and size selection were performed with cutadapt v4.1 (-u 1 -a TGGAATTCTCGGGTGCCAAGG --minimum-length 19 to --maximum-length 25) (Martin 2011). Processed reads were mapped to the Kronos v1.1 genome for *MIR* and *PHAS* annotation.

The *MIR* loci were annotated by aligning precursor sequences from Svevo v1 to the Kronos assembly using blastn v2.16.0 (Camacho et al. 2009), retaining the best hit per precursor with the highest sequence identity on the syntenic chromosome. Mature miRNA sequences were aligned with blastn-short (-word_size 15 -perc_identity 100 -no_greedy -ungapped -dust no -outfmt 6). Alignments were retained if there were no mismatches in the central region (positions 5–18 of the query) and ≤ 4 total mismatches. To estimate miRNA abundance and reassess annotation confidence, ShortStack v4.1.1 was used (--dicermin 20 --dicermax 24 --mmap u --mincov 1.0 --pad 75 --threads 20 --dn_mirna -- locifile and --known_miRNAs miRBasePlantMirnas.fa) (Axtell 2013; Johnson et al. 2016). Loci predicted as siRNA rather than miRNA were designated as low confidence. Normalized expression was calculated for each locus (reads per million; RPM).

*PHAS* loci were identified with ShortStack v3.8.5 (Axtell 2013; Johnson et al. 2016), using phase scores of ≥40 and RPM abundance of ≥2.0 as thresholds. Loci producing non-phased 24-nt siRNAs were classified as putative heterochromatic siRNA (hc-siRNA) loci. *PHAS* loci were categorized by phasiRNA length (21-nt or 24-nt) and peak accumulation stage (premeiotic, meiotic, or postmeiotic). Overlaps between protein-coding genes (v2.1) and sRNA loci were examined with Bedtools v2.31.1 with the intersect command and a minimum overlap fraction (*-*f) of 0.75 (Quinlan and Hall 2010).

### Reciprocal placement of *PHAS* loci between Svevo and Kronos

To assess conservation of *PHAS* loci, we performed reciprocal sequence similarity searches between Svevo and Kronos genomes using blastn v2.16.0 (Camacho et al. 2009). For each precursor, only the top hit was retained, with preference given to matches on the syntenic chromosome. If no syntenic hit was identified, the single best match hit was kept.

### EC data analysis

EC sequencing data from 1,479 samples, designed for 1,439 Kronos lines, were downloaded from PRJNA258539 (Krasileva et al. 2017). Adapter trimming and quality filtering were performed using fastp v0.23.4 (Chen et al. 2018). Filtered reads were aligned to the Kronos genome using bwa aln v0.7.18-r1243-dirty (Li 2013). The resulting alignments were sorted by samtools v1.20 (Li et al. 2009), and duplicates were removed using picard v3.0.0. Processed alignments were analyzed using the MAPS pipeline (Henry et al. 2014), organized in batches of approximately 24 each, consistent with the structure of mutant library generation (Krasileva et al. 2017).

For a mutation to be considered valid, a base had to be initially covered by at least one mapped read with mapping quality greater than 20 in at least (N - 4) samples, where N corresponds to the number of alignments in a batch. Multi-mapped reads were re-examined, and their mapping qualities were corrected using the internal multi-mapped read recovery pipeline within MAPS, after which mutations were reassessed to meet these criteria. Reliable mutations required additional evaluation using HetMC and HomMC. Consistently with previous studies (Krasileva et al. 2017; Zhang et al. 2023b), the HetMinPer parameter was fixed to 15, and four HetMC and HomMC pairs were set from the lowest to highest stringency: (3, 2), (4, 3), (5, 3) and (6, 4). Mutations passing the initial filter and detected from uniquely mapped reads in genomic regions without residual heterogeneity were further evaluated by these thresholds. The least stringent HetMC and HomMC thresholds yielding ≥98%, ≥97%, and ≥95% EMS mutation rates were used to call high, medium and low confidence mutations, respectively. If none of these thresholds achieved ≥98% EMS mutation rates, high, medium and low confidence mutations were called using a predefined order of HetMC and HomMC as in the previous study (Krasileva et al. 2017): (5, 3), (4, 3) and (3, 2). For all detected mutations, variant effects were predicted using snpEff v5.2a on the annotation v2.1

### PC data analysis

PC sequencing data for 1,641 samples were obtained either directly from the authors of the previous study or downloaded from PRJNA1218005 (Zhang et al. 2023b). Sequencing reads were trimmed with trimmomatic v0.39 (Bolger et al. 2014), and alignments were generated using the identical workflow described for EC data. Following the BioSample organization, about 48 alignments were processed per batch, with the exception that PC09 samples were excluded due to quality issues, and PC7 and PC8 samples were processed as a single batch. The required library size was adjusted to (0.8 x N), where N corresponds to the number of alignments in the batch, following the previous study (Zhang et al. 2023b). In total, 1,556 Kronos mutants were processed. Downstream mutation retrieval using the MAPS pipeline followed the same workflow as for EC data.

### Mutation discovery using GATK

To provide comprehensive variant resources, mutations were identified from Kronos EC and PC datasets using GATK HaplotypeCaller v4.5.0 (McKenna et al. 2010). Trimmed sequencing reads were aligned to the Kronos genome using bwa aln v0.7.18-r1243-dirty, and duplicate reads were marked using picard v3.0.0 prior to HaplotypeCaller (Li 2013).

### Gene expression analyses

Raw RNA-seq datasets were downloaded from NCBI (Table S32), and adapters and low-quality reads were removed using fastp v0.24.0 (-q 20 --length_required 50) (Chen et al. 2018). For quantification, we used the Kronos v2.1 annotation together with the reference genome to generate a decoy-aware index. Gene-level abundances were then estimated with Salmon v1.10.3 (--validateMappings) (Patro et al. 2017).

For each BioProject, gene-level counts were imported into R v4.3.2, and replicate consistency was assessed with PCA analyses (Ihaka and Gentleman 1996). Outlier samples were excluded to ensure stability of cross-study comparison. For the remaining samples, counts scaled by effective transcript length (lengthScaledTPM) were normalized using the trimmed mean of M values (TMM) methods in EdgeR v4.0.16 and converted into CPM (Robinson et al. 2010). Within each sample, CPM values were ranked against the full transcriptome, and percentiles were calculated from 0 (lowest expression) to 1 (highest expression). Biological replicates were aggregated by calculating median percentiles per condition. For tissue-level comparisons, samples were grouped into leaf, endosperm, and meristem, with only wild-type meristem samples included. Median percentile ranks were then calculated for each NLR within each tissue.

## Data availability

The sequencing data has been deposited in the National Center for Biology Information under BioProject PRJNA1213727. All datasets can be accessed through Zenodo: https://zenodo.org/records/10215402 and https://zenodo.org/records/15399687. To identify datasets of interest, please refer to Table S1. Our genomic databases are hosted in the GrainGenes (https://graingenes.org/GG3/).

## Code availability

Computational pipelines and custom scripts are available at https://github.com/s-kyungyong/Kronos. Scripts and data for hvNLR analyses are available at https://github.com/daniilprigozhin/Wheat_NLRome.

## Contributions

K.V.K. conceived research. K.S. and K.V.K. conceptualized the project. R.K. extracted DNAs for PacBio sequencing. D.P. developed the improved hvNLR identification pipeline and performed hvNLR predictions. C.L. prepared proximity libraries and coordinated sequencing. T.H.C.R., S.B, J.W.H. and B.C.M. conducted *MIR* and *PHAS* analyses. K.S. curated all datasets and performed the full suite of computational analyses, with the exception of *MIR* and *PHAS* analyses. M.T. assisted with syntenic analyses and data curation. K.S. wrote the manuscript with input from K.V.K and C.L.

## Acknowledgements

We thank Dr. Junli Zhang, Dr. German F Burguener and Dr. Jorge Dubcovsky for their suggestions to improve genome annotations and assistance with PC dataset transfer. K.S. used ChatGPT 4o and 5 to assist with coding. This research relied on the Savio computational cluster resource provided by the Berkeley Research Computing program at the University of California. K.S. is supported by the Berkeley BioEnginuity Fellowship. The project received support from the Department of Agriculture-National Institute of Food and Agriculture (2021-67013-35726) to KVK. Work on miRNA and phasiRNA annotation in the Meyers lab is supported by US National Science Foundation award DBI-2450802. The Berkeley Center for Structural Biology is supported by the Howard Hughes Medical Institute, Participating Research Team members, and by the National Institutes of Health, National Institute of General Medical Sciences grant P30 GM124169.

