## Supplemental Figures for "The Annotated Blueprint: Integrated Functional Genomic Resources for a model Tetraploid Wheat *Triticum turgidum* cv. Kronos"

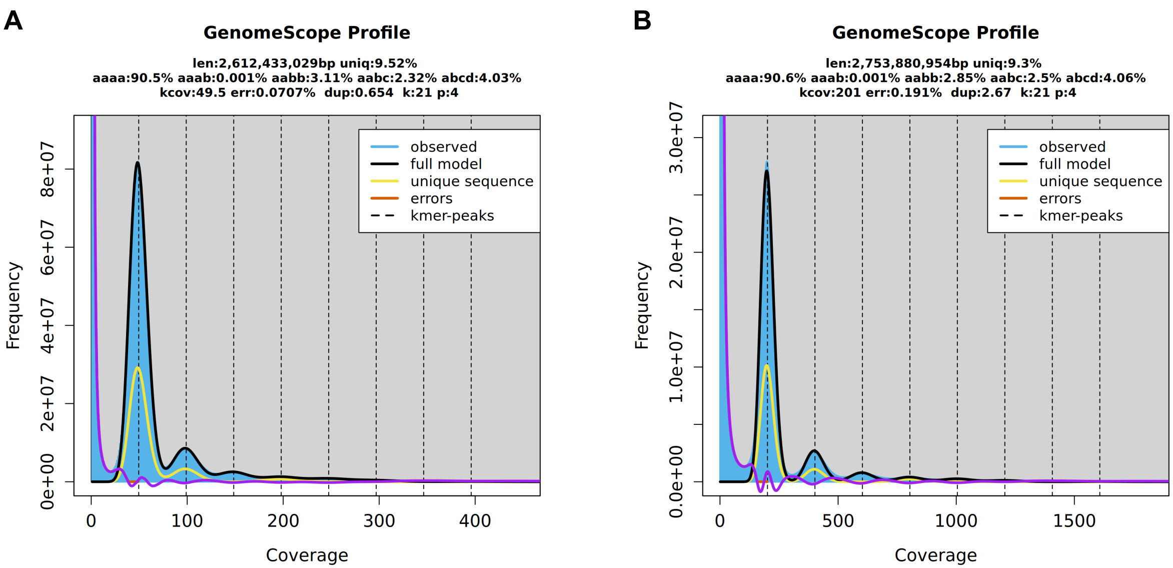


**Figure S1. Comparative genome profiling of Kronos and Svevo using GenomeScope**

GenomeScope k-mer profile plots for **(A)** Kronos, generated from PacBio HiFi reads in this study, and **(B)** Svevo, generated from publicly available Illumina paired-end reads (PRJEB22687). The profiles illustrate genome size, ploidy, and heterozygosity estimates derived from sequencing data.

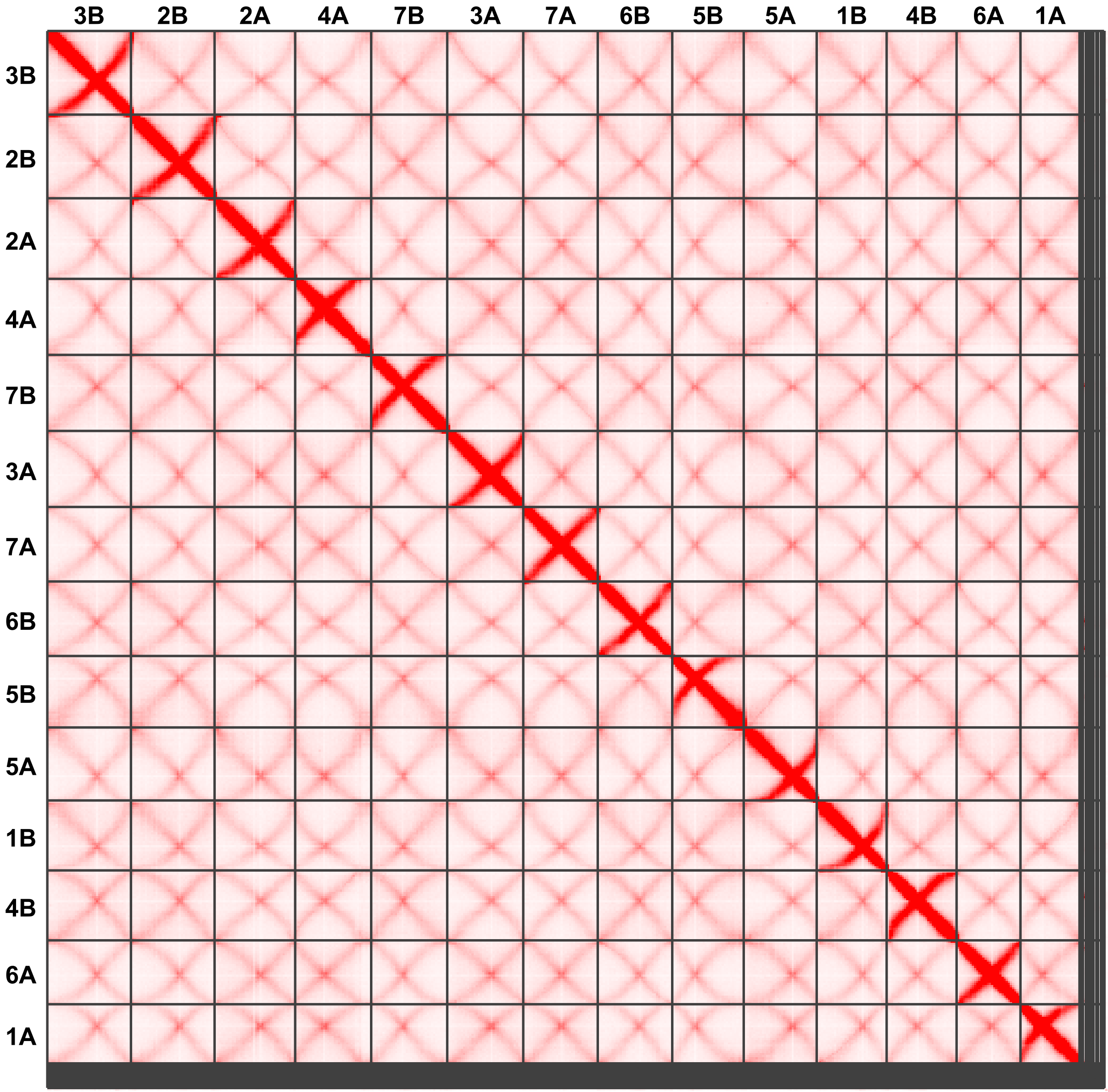


**Figure S2. Hi-C chromatin contact maps of Kronos chromosomes (5 kb resolution)**

Hi-C contact maps of Kronos chromosomes at a resolution of 5 kb, visualizing chromatin interactions along the whole chromosomes. This interaction maps originally display scaffolds from the largest to smallest, which were later renamed in consistency with the Chinese Spring reference genome.The orientation follows the Kronos reference genome v1.0. Note that In v1.1, the orientations of 1B, 2A, 2B, 3A, 3B, 5A, 6A and 6B were flipped to match the Chinese Spring reference genome. The intensity of red coloration indicates the frequency of chromatin interactions, with stronger interactions along the diagonal, reflecting intra-chromosomal contacts.

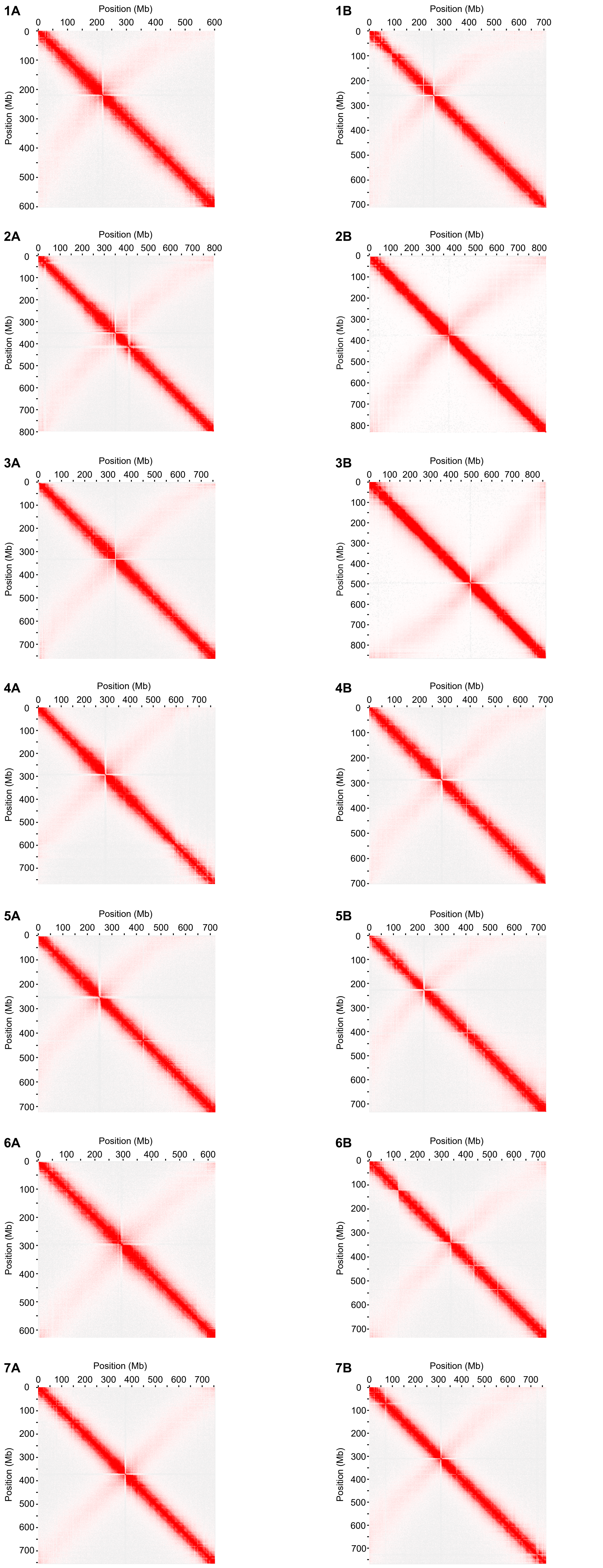


**Figure S3. Hi-C chromatin contact maps of Kronos chromosomes (2.5 Mb resolution)**

Hi-C contact maps of Kronos chromosomes at a resolution of 2.5 Mb, visualizing chromatin interactions along the chromosome length. The orientation of chromosomes is consistent with the Kronos reference genome v1.1. The x- and y-axes represent genomic coordinates (Mb). The intensity of red coloration indicates the frequency of chromatin interactions, with stronger interactions along the diagonal, reflecting intra-chromosomal contacts.

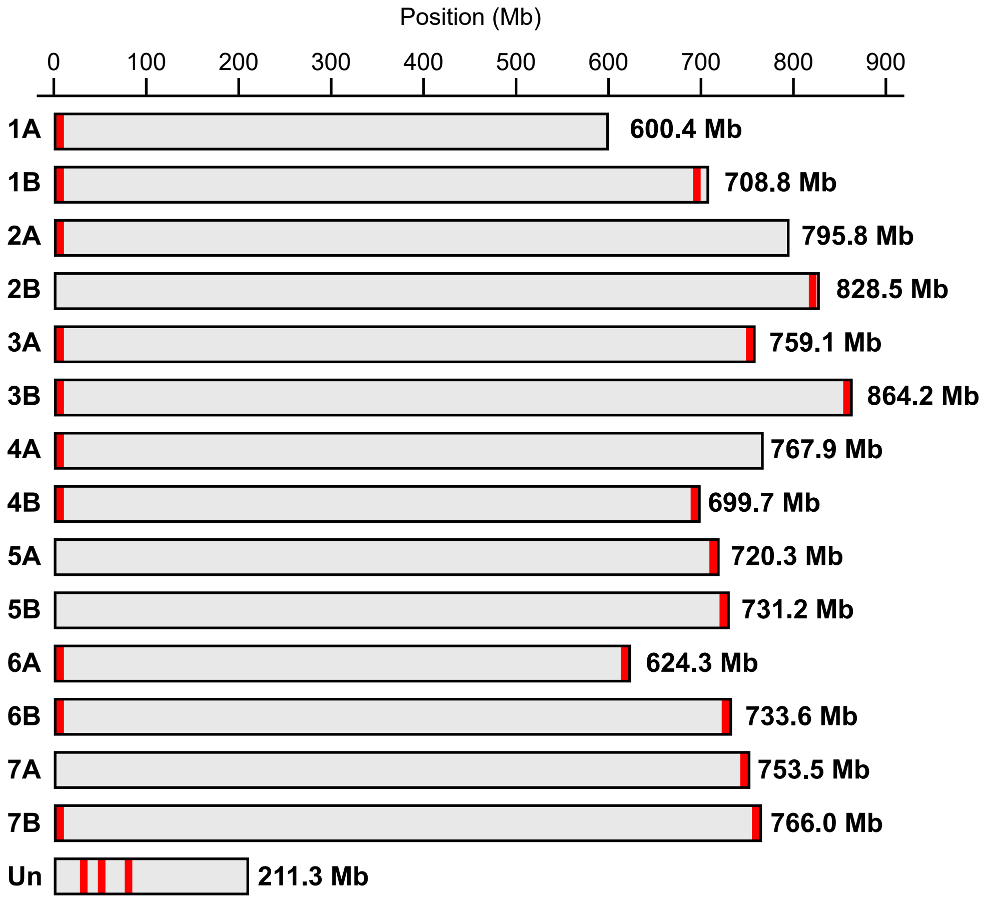


**Figure S4. Chromosome sizes and telomeric repeat distribution in the Kronos genome**

Chromosome sizes of the Kronos reference genome and the distribution of telomeric repeats. Telomeric repeat regions were identified using tidk, based on the occurrence of the AAACCCT motif characteristic of Poales. A minimum threshold of 100 repeats within a 10,000 bp window was required for visualization.

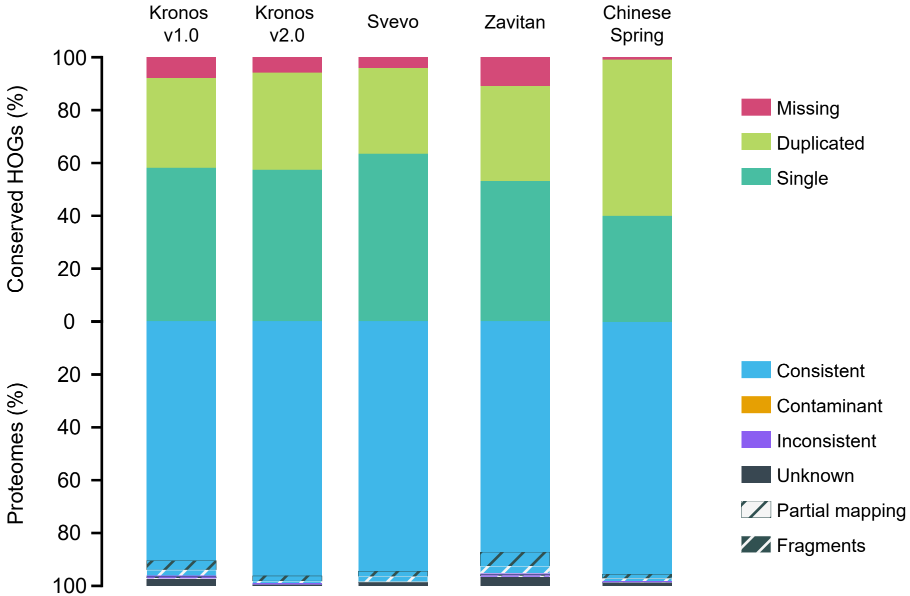


**Figure S5. Proteome quality assessment of Kronos and related wheat genomes using OMArk**

Quality assessment of annotated proteins using OMArk. Evaluation includes (i) consistency of 10,157 conserved hierarchical orthologous groups (HOGs) in Triticinae and (ii) proteome-wide assessment against known gene families. Results are shown for Kronos versions and related wheat genomes.

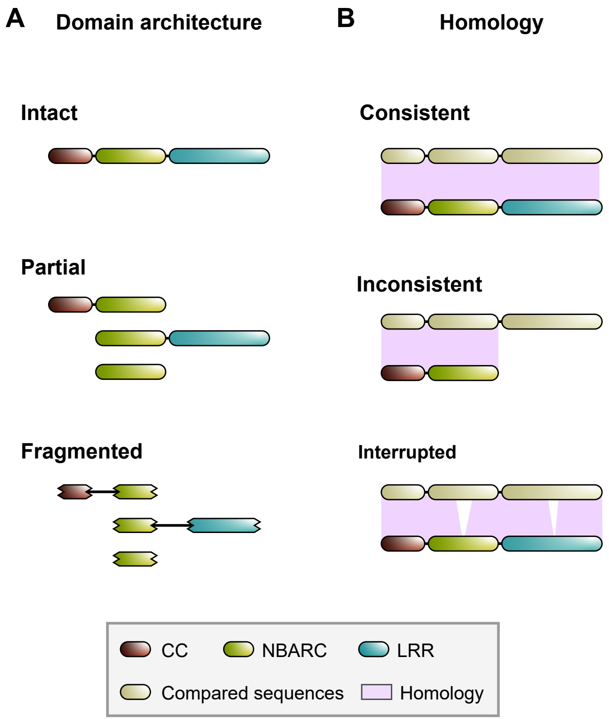


**Figure S6. Manual curation of NLRs based on domain architecture and homology**

All manually curated NLRs were annotated with two labels: **(A)** domain architecture and **(B)** sequence homology. **(A)** Domain architecture classification. **Intact**: Contains a canonical domain structure with an N-terminal coiled-coil (CC) domain, a central NB-ARC domain, and a C-terminal leucine-rich repeat (LRR) domain. **Partial**: Contains a detectable NB-ARC domain but lacks either the CC or LRR domain. **Fragmented**: Composed of incomplete or truncated domain remnants, lacking clear canonical structure. **(B)** Sequence homology classification. **Consistent**: displays global or near-complete homology with NLR sequences in the NCBI non-redundant database. **Inconsistent**: Lacks major domain regions or shows substantial deviation from known reference sequences. **Interrupted**: Contains internal gaps due to frame disruptions. These regions are bypassed via unsupported splice junctions during transcript model generation.

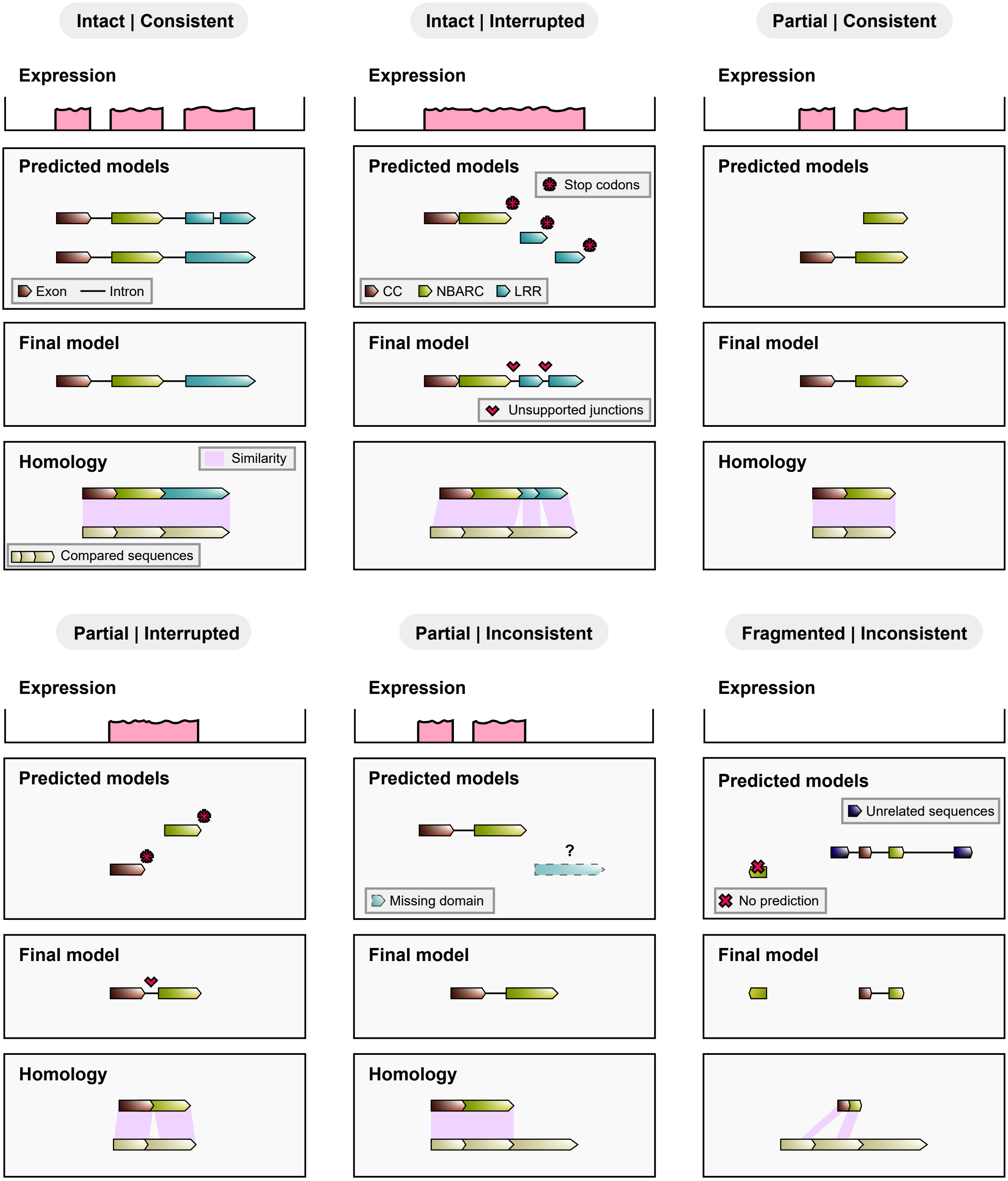


**Figure S7. Major annotation categories of curated NLRs**

Manually curated NLRs were classified into major categories based on domain integrity, sequence homology, gene model quality, and transcriptional support. Although these categories often correspond to shared characteristics in gene prediction and expression, they are not strictly mutually exclusive. Ambiguous cases were assigned to the most representative category based on available evidence. **Intact | Consistent**: exon-intron junctions are well-supported by transcriptome data and/or homologous sequences with global alignments. **Intact | Interrupted**: canonical gene structures are disrupted by one or more mutations introducing premature stop codons or frameshift mutations. Gene predictors may insert unsupported introns to bypass these mutations or produce multiple fragmented gene models. When transcriptomic data is available, gene expression patterns may not align with predicted gene structures. Arbitrary splicing sites are introduced during manual curation to retain intact domain architecture, and therefore, equence gaps appear in homolog alignments. Additional labels, such as "**1 | NB-ARC**," indicate the estimated number and location of mutations that interrupted the NLR structure. **Partial | Consistent**: NLRs lack either an N-terminal coiled-coil (CC) domain or a C-terminal leucine-rich repeat (LRR) domain. Nevertheless, exon-intron junctions are well-supported by transcriptome data and/or homologous sequences consistently found in multiple wheat species. **Partial | Interrupted**: similar to the previous class, but with mutations interrupting the gene structure. Arbitrary splicing sites are introduced as for Intact|Interrupted. **Partial | Inconsistent**: NLRs without intact domain architecture are annotated, but close homologs contain the missing domains, suggesting their potential loss in the annotated sequences. **Fragmented | Inconsistent**: All NLRs labeled as fragmented are inconsistent and likely pseudogenes. Gene predictors either fail to annotate models or annotate models with fragmented domains. These models may lack start and/or stop codons and include unreliable splicing sites.

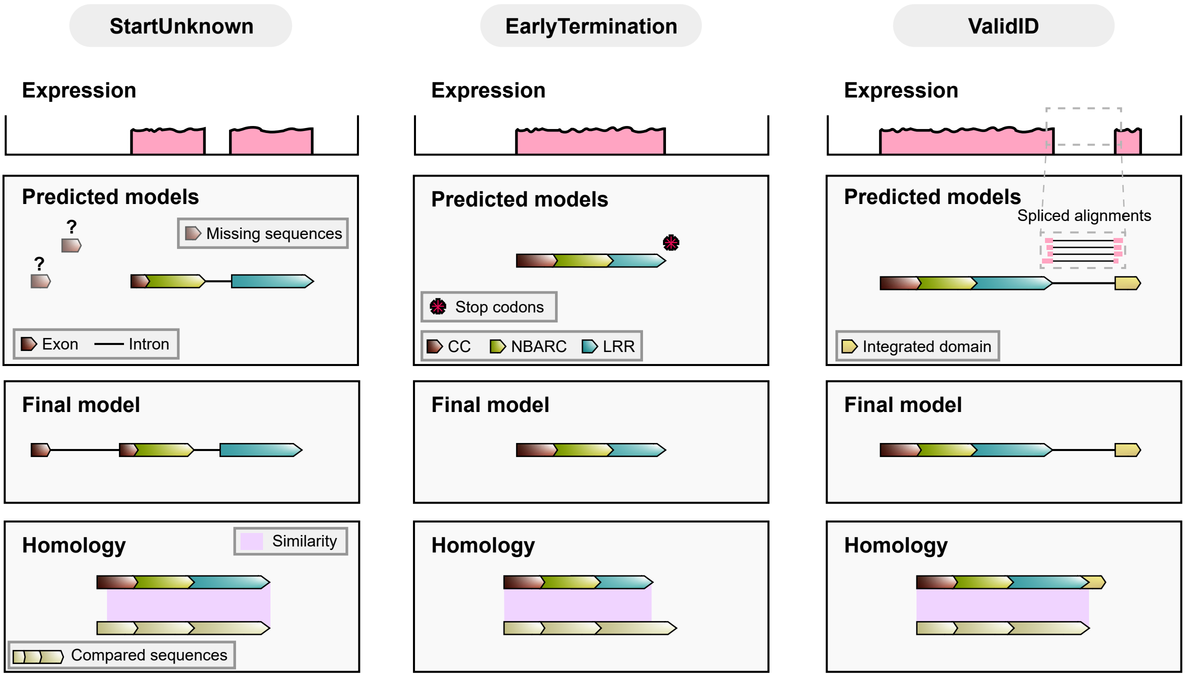


**Figure S8. Additional annotation categories for manually curated NLRs**

In addition to primary classifications, some NLRs were assigned secondary labels to highlight specific structural or functional features identified during curation: **StartUnknown**: Supported gene models lack a start codon, but the initial exons or sequences that contain a start codon could not be confidently identified. **EarlyTermination**: The gene model, particularly leucine-rich repeat domain, terminates earlier than expected based on compared sequences in the database. This early termination may or may not be supported by transcriptome data. **ValidID**: Curated NLRs contain integrated domains (IDs), and the splicing junctions between canonical NLR domains and IDs are supported by transcriptome data. Without the transcriptomic evidence, the gene is assigned to the **PutativeID** class.

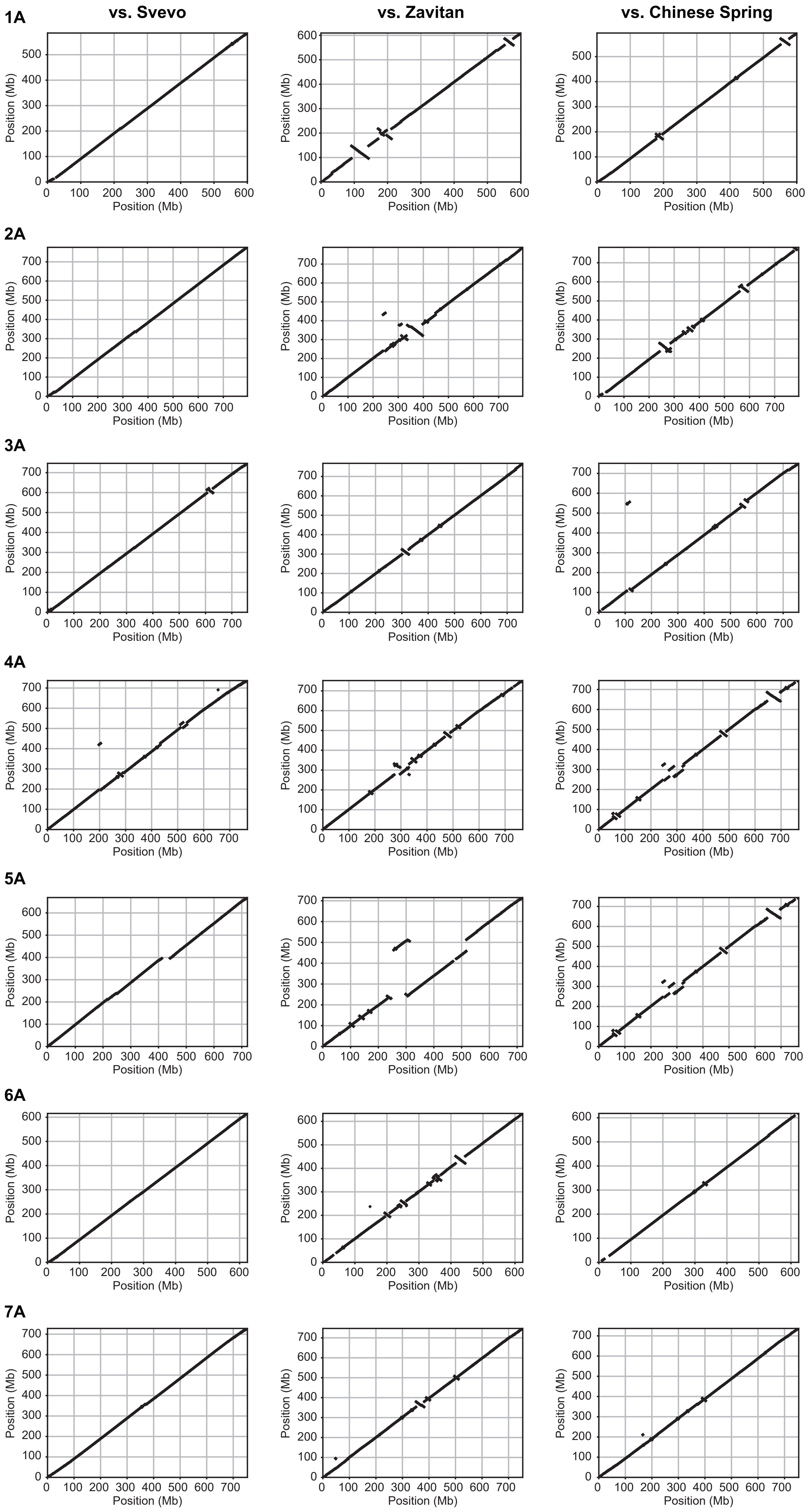


**Figure S9. Chromosome-scale synteny of the A subgenome between Kronos and wheat reference genomes**

Dot plots show chromosome-wide syntenic relationships between Kronos (x-axis) and three reference genomes (y-axis): Svevo (left), Zavitan (middle), and Chinese Spring (right).

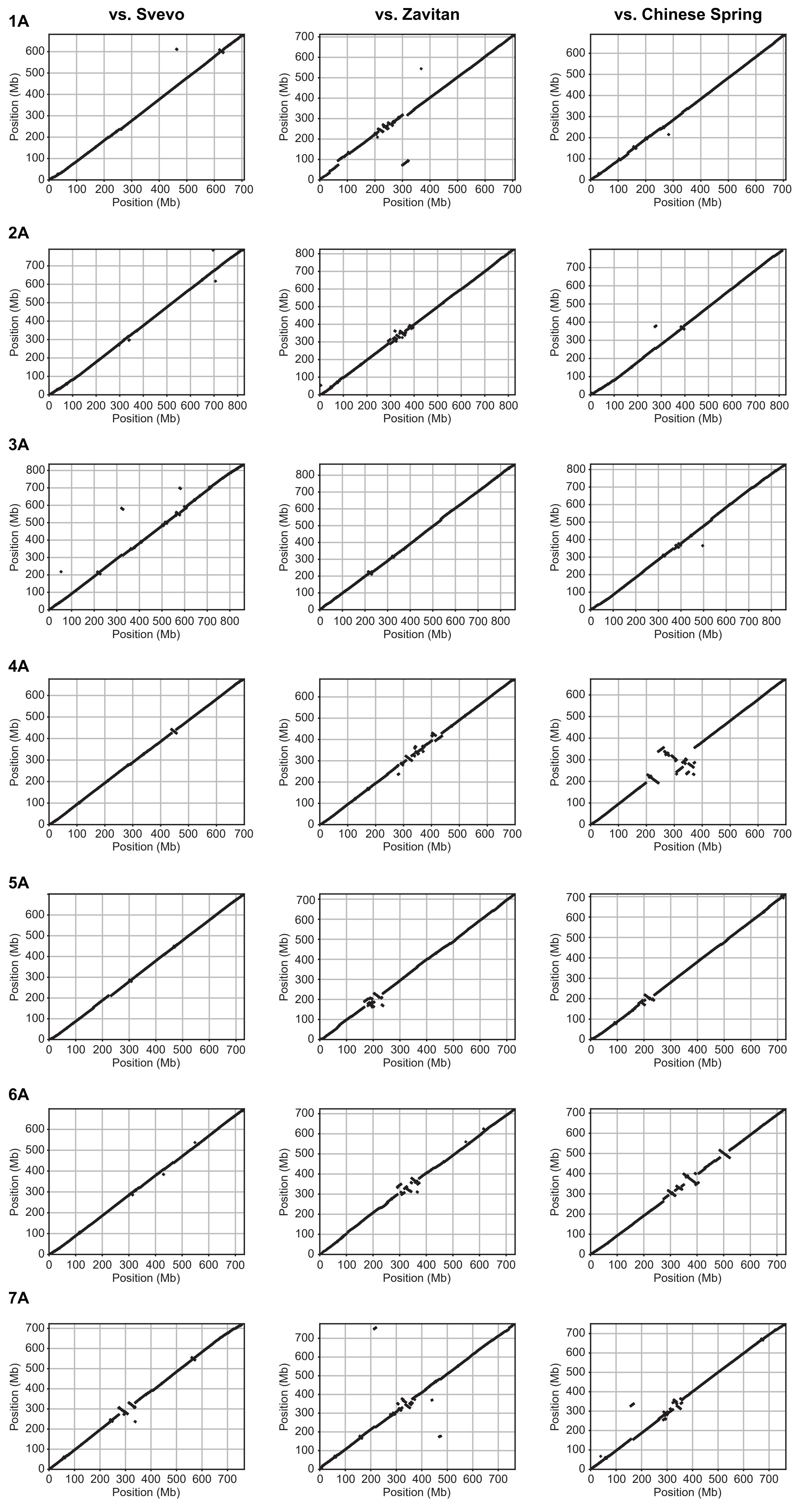


**Figure S10. Chromosome-scale synteny of the B subgenome between Kronos and wheat reference genomes**

Dot plots show chromosome-wide syntenic relationships between Kronos (x-axis) and three reference genomes (y-axis): Svevo (left), Zavitan (middle), and Chinese Spring (right).

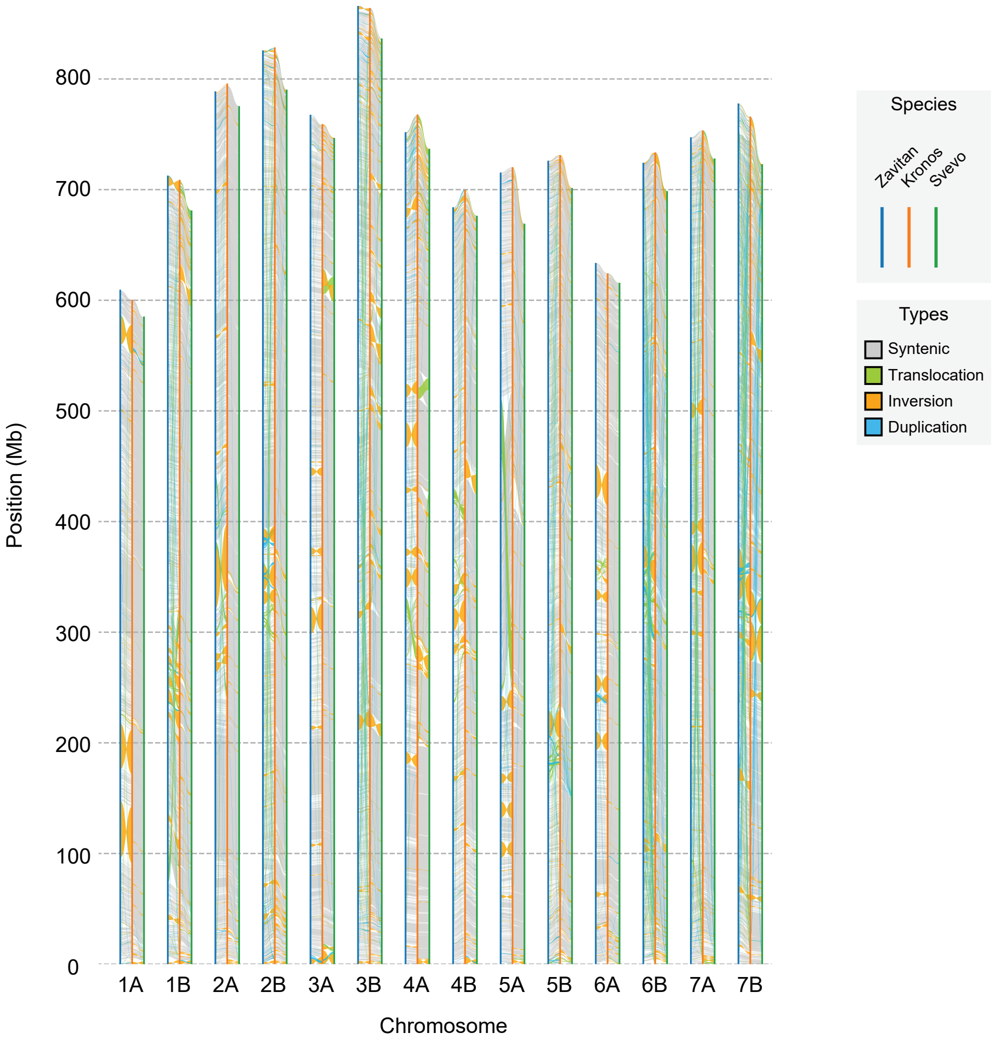


**Figure S11. Genome-wide synteny and structural rearrangements identified with SyRI**

Chromosome-scale comparisons of Kronos (orange), Zavitan (blue), and Svevo (green) were performed using SyRI. Four types of structural features were identified: syntenic regions (gray), translocations (green), inversions (orage), and duplications (blue). The vertical axis indicates chromosomal positions. The plot highlights both conserved syntenic blocks and lineage-specific structural variations among the three cultivars.

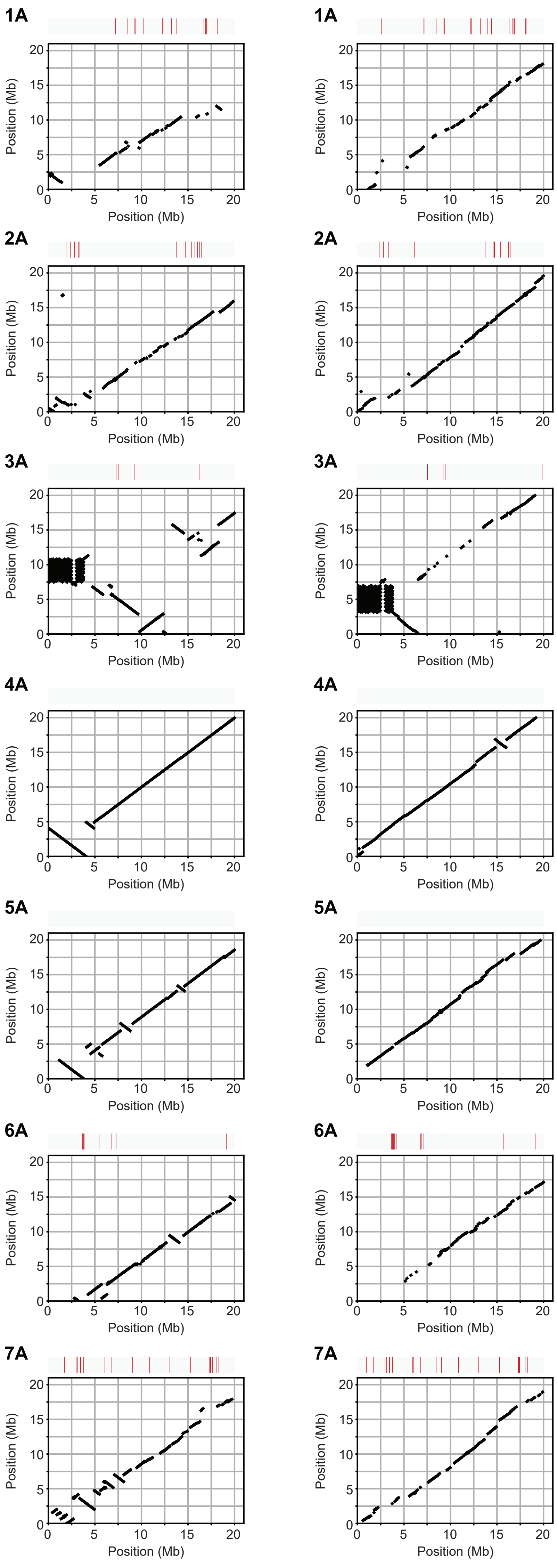


**Figure S12. Chromosome head synteny of the A subgenome in Kronos compared with Svevo and Zavitan**

Dot plots show local syntenic relationships across the first 20 mega base pairs (Mb) of each A subgenome chromosome. The x-axis indicates the positions in the Kronos genome, and the y-axis shows the corresponding positions in the compared reference genomes: Svevo (left) and Zavitan (right). The tracks above dot plots indicates the positions of manually curated reliable NLRs in Kronos (presence: red and absence: grey).

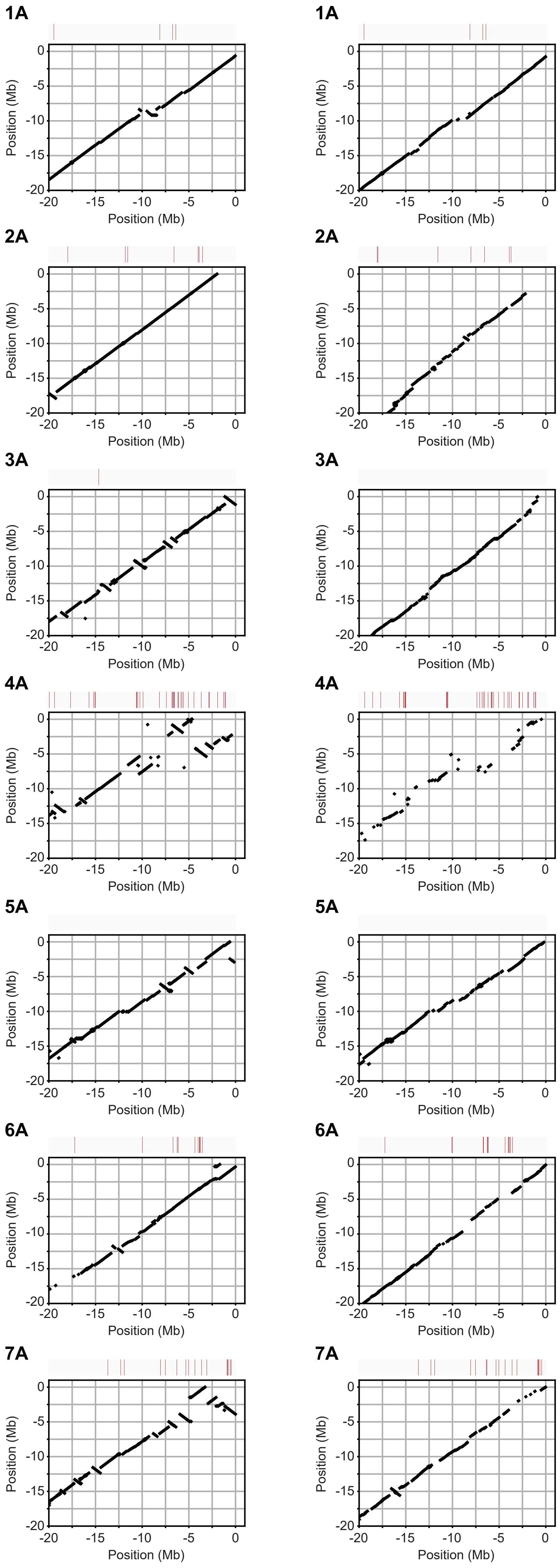


**Figure S13. Chromosome tail synteny of the A subgenome in Kronos compared with Svevo and Zavitan**

Dot plots show local syntenic relationships across the last 20 mega base pairs (Mb) of each A subgenome chromosome. The x-axis indicates the positions in the Kronos genome, and the y-axis shows the corresponding positions in the compared reference genomes: Svevo (left) and Zavitan (right). The position 0 marks the end of each chromosome, and negative values indicate relative distance upstream from the end. The tracks above dot plots indicates the positions of manually curated reliable NLRs in Kronos (presence: red and absence: grey).

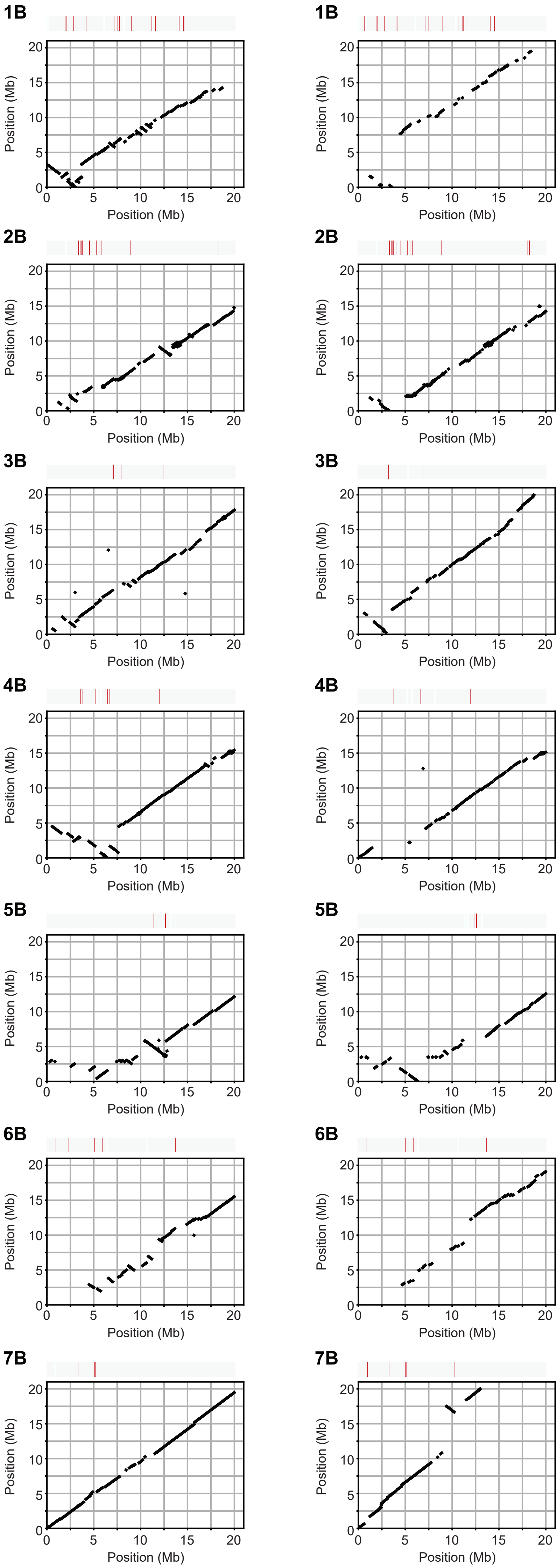


**Figure S14. Chromosome head synteny of the B subgenome in Kronos compared with Svevo and Zavitan**

Dot plots show local syntenic relationships across the first 20 mega base pairs (Mb) of each B subgenome chromosome. The x-axis indicates the positions in the Kronos genome, and the y-axis shows the corresponding positions in the compared reference genomes: Svevo (left) and Zavitan (right). The tracks above dot plots indicates the positions of manually curated reliable NLRs in Kronos (presence: red and absence: grey).

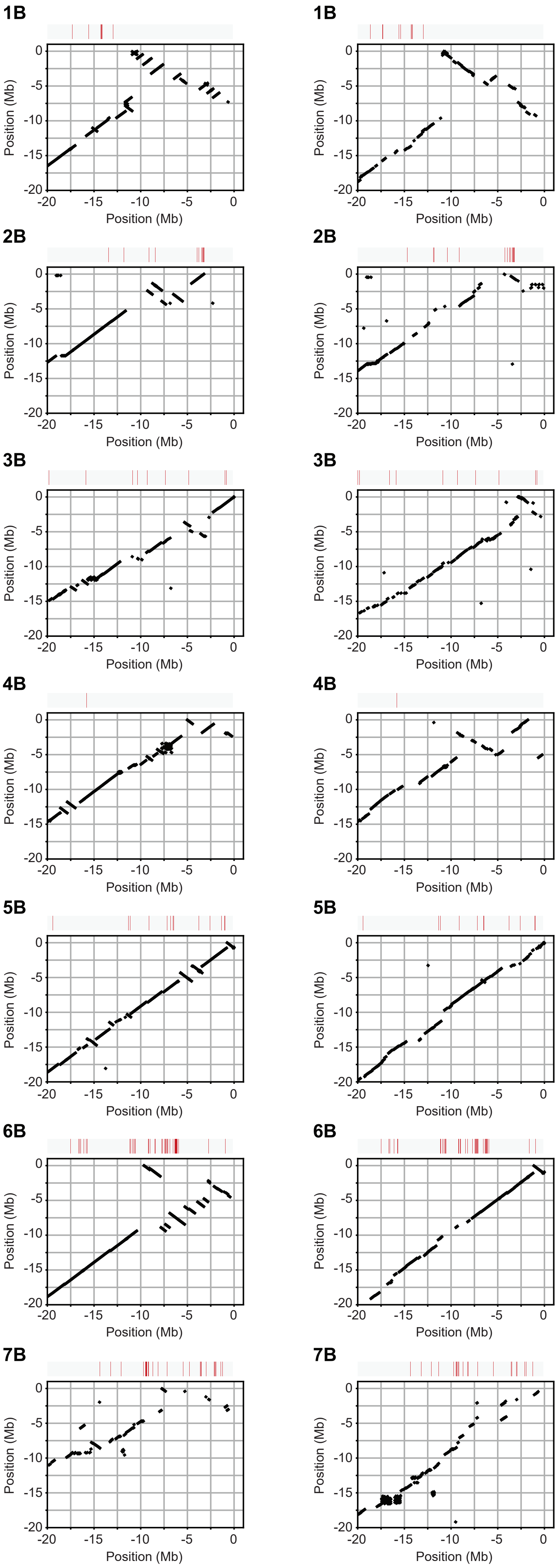


**Figure S15. Chromosome tail synteny of the A subgenome in Kronos compared with Svevo and Zavitan**

Dot plots show local syntenic relationships across the last 20 mega base pairs (Mb) of each B subgenome chromosome. The x-axis indicates the positions in the Kronos genome, and the y-axis shows the corresponding positions in the compared reference genomes: Svevo (left) and Zavitan (right). The position 0 marks the end of each chromosome, and negative values indicate relative distance upstream from the end. The tracks above dot plots indicates the positions of manually curated reliable NLRs in Kronos (presence: red and absence: grey).

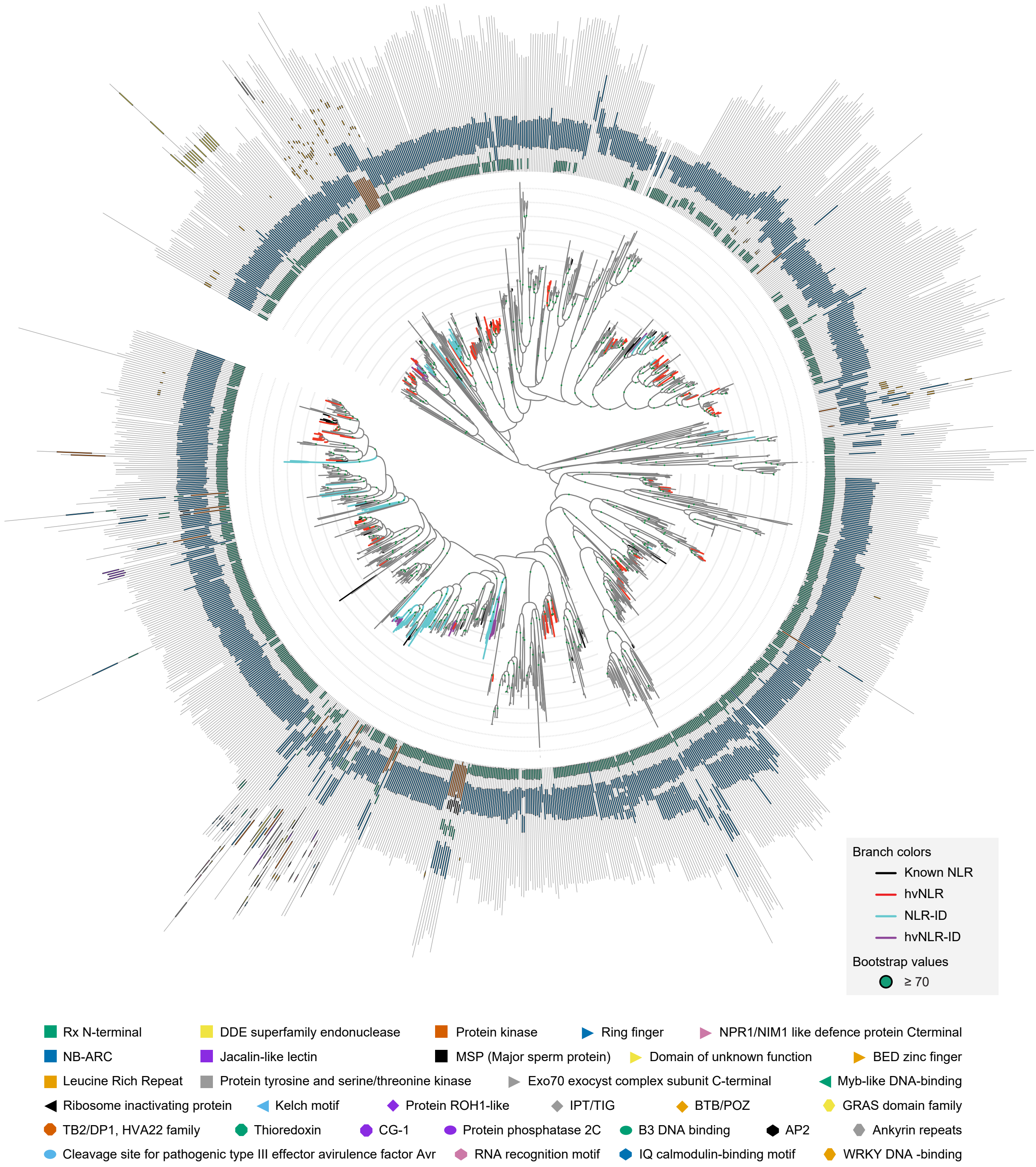


**Figure S16. Domain architectures and phylogenetic distribution of Kronos NLR-IDs**

Maximum-likelihood phylogeny inferred from NB-ARC domains. Branch colors indicate: black—cloned, functionally validated NLRs; red—Kronos highly variable NLRs (hvNLRs); sky blue—Kronos NLR with PFAM-annotated integrated domains (NLR-ID); purple—Kronos hvNLR-IDs. Bars around the circular tree indicate domain architecture, with functional categories color-coded as shown in the legend. Bootstrap supports ≥ 70 are shown as green circles at the corresponding nodes. For a detailed view of gene identifiers and domain labels, please refer to the accompanying datasets.

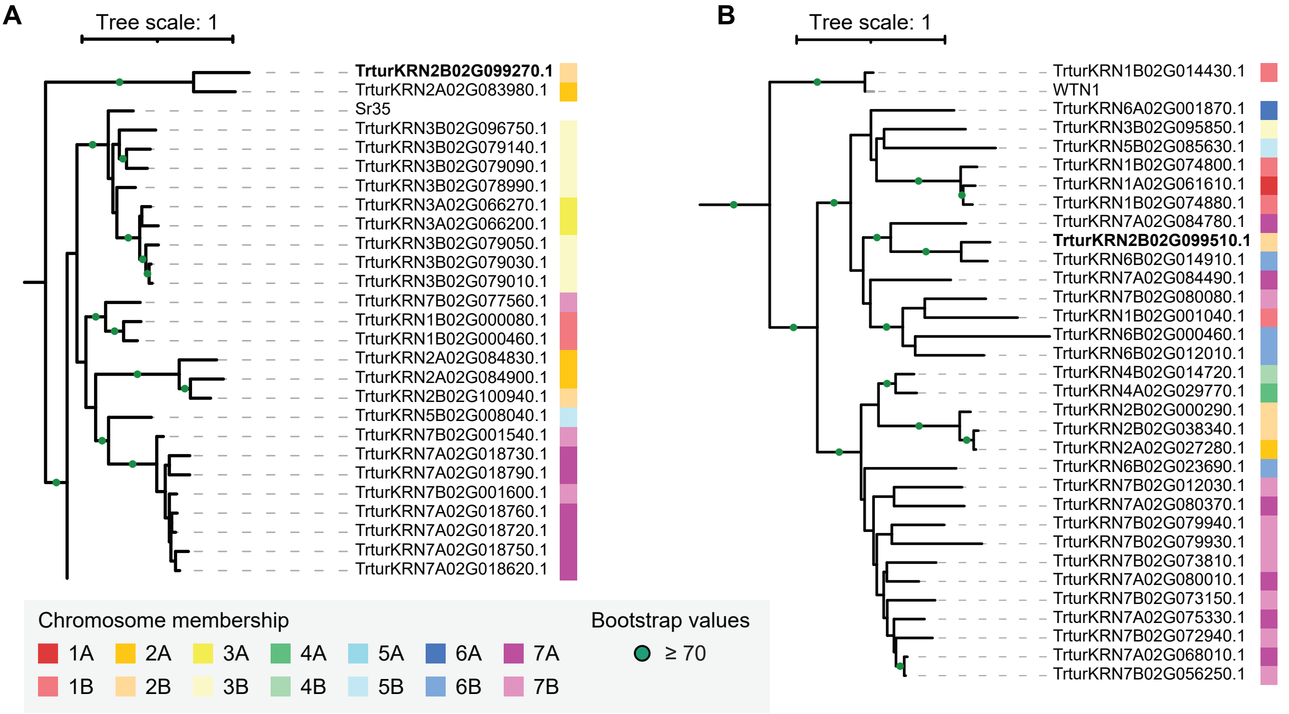


**Figure S17. Phylogenetic placement of Kronos NLRs TrturKRN2B02G099270 and TrturKRN2B02G099510**

Maximum-likelihood phylogenetic trees display the relationships of TrturKRN2B02G099270 and TrturKRN2B02G099510 with homologous NLRs  Terminal labels indicate protein names. Colored squares to the right of each tip denote chromosome membership. Nodes supported with bootstrap values ≥70 are marked with green circles.

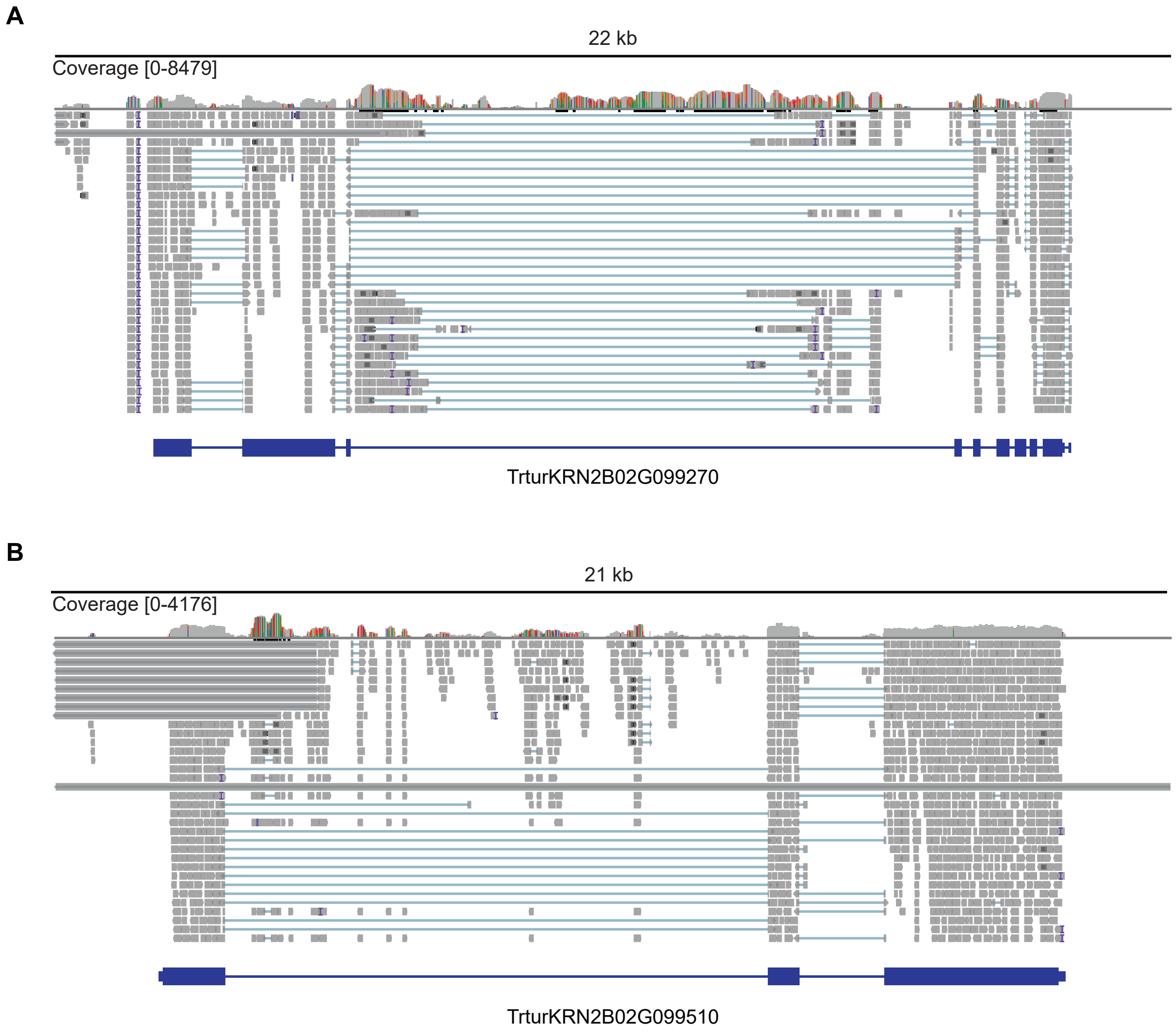


**Figure S18. Read coverage and alignment tracks for TrturKRN2B02G099270 and TrturKRN2B02G099510**

**(A)** TrturKRN2B02G099270 and **(B)** TrturKRN2B02G099510. The upper track shows coverage depth, while the lower track displays individual read alignments in grey. Gene models are shown at the bottom, with exons highlighted in dark blue.

**A**

\>TrturKRN2B02G099510.1
MPPDLTLHSLLNPISVCSVDAQFPSLASTHHLLAAPQTKTTQGGRQSGRATMANFALGLAKTAVEGTLSR
VLSAIDEENMLKVAVQQDLVFITGEFQMMQSFLNIASKERARNEVVRTWVRQLRDLAFDVEDCVEFVIHL
DNDSTRNWVWRLVHSCMAPPRTRVLDDTIAELKQLKARVEEVSKRNTRYNLISDSSSNTVSQSPAGQLAT
AGPSTIDILMELWVANGKYLGFCNLQKLITRESNDLQVISLWGSAGSDIEGAQIINKAYCDPEISRIFKV
RAWVKLMHPFSHDEFLKSLQTQLQASCRCCEAVHSKGESSENDPMKQLTEQRYLIILDGIPSVVEWDIIR
MHLPENNNGSRIVVSTKQLGDAIFFTGKHYRVSVLREFSGGQYICAFYKKASDCYVYEDNEKASRSTFLS
KRMQKRVPLSSQKYVLINIMNNWDRTGVTSVWGIAGVGKSTLVGDRYYQRMQDCSLVGKFAWVDVPDPFD
LVEFSRLLLLDFHRCDLQAMEAAAIHLMQGQDPIQGCREILRQDGYLVIIVGLQSTHDWDLIETAFLSET
TTRANIVVITTEESIAKHSVKQQVDQMINIKCIEDDEAYDLFKEIIDHKELAILKEKKVDLAKLVAKCGG
LLKVVSSIGQVSGSLLEHMNDDFMVKLETYPEFHSLSGLLSWMDSYFDACSDLVKPCIFYLSVFPASSHI
RRCRLLRRWIAEGYSRDTADCTAKENAQKLFSELIGLSIIQQPRNNDICQVNCFFREYIISRPMVDNLVF
ALDGRCSPNTPRTGQHLTIMRNWHRDQIVFERIDLTRLRSLTIFGTWCSFFISDKMKHLRVLDLEGMIAI
TNGDIEQIFVKLLSLKFISLRGCKNISRLPDTLGGLRQLQTLDVRFTSIVMLPSAIIKLQKLQYIRAGTD
QDWTSASLEDGDGITS**QLPATTSPEQERLDSIISASQPTVRTSPEIGIAESPAHEATSPEQEGLDSIIYA**
**SQPTVPTSFEIGIAKSPAHAATSPEQEGLDSIISASQPTVPTSPEIGIAESPAHAATSPEKEGLDSIISA**
**SQPTVPTSFEIGIAQSPAHEATSPEEKGLDRIISASQPTVPTSPEIGIAESPAHAATSPEQECLDRIISA**
**SQPTVPTSPEIGIAKSPAHAATSPEQEGLDSIISASQPTVPTSPEIGIAESPTHAATSPQQEVLDRIISA**
**SQPTVPTSLEIGIAESPAHEATSPEQEGLESIISASQPTVPTSPEIGIPESPAHATTSPEQEGIDSIISA**
**SQPTVSTSPEIGIAKSPAHAATSPEEEGLDSIISASQPTVRTSPEIGIAESPAHAATSPEQEGLDSIISA**
**RQPTMPKSPEIGIAESPAHEATSPEQEGLDSIISASQPIVPTSPEIGIAESPAHAATSPEQEGLDSIISA**
**SQPTVPTSPEIGIAESPSQAE**RSAKAPNMPIKPKVGWASKLSCNSVPRLPNFPPVTWSLFGSEREKEAQY
YSAGVEVPARIEKLAVLHTIGVVNINSSGGEAFLKKLTQLAHLVKLRVCGINKKNWQRLCSALSIHGHLE
SLSVRFDENCLDDSFKPPKTLKSLKLYGPIRKLPDTIKVLDNLKKFDLEMTITGSEDMHVFMKDKLPRKD
ILNRLCIKVQGVQKVNFGSRELGHHSLFKPRVVKIDCSSKLHVTFGYDGTSSNVEVLIIQCSKGSSFRVT
GENELLVMHRLKTVWLKGSYNEEQQLHLKELVDRHRNKAIVLKLEPL

**B**

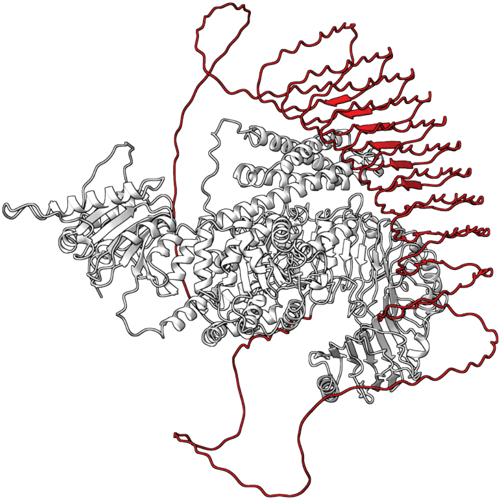

**Figure S19. Sequence composition and predicted structure of TrturKRN2B02G099510**

**(A)** Amino acid sequences of TtrurKRN2B02G099510, with simple repeat motifs highlighted in red (residues 927–1421). **(B)** Predicted structure of TtrurKRN2B02G099510 from AlphaFold, displaying the spatial distribution of the simple repeat regions in red.

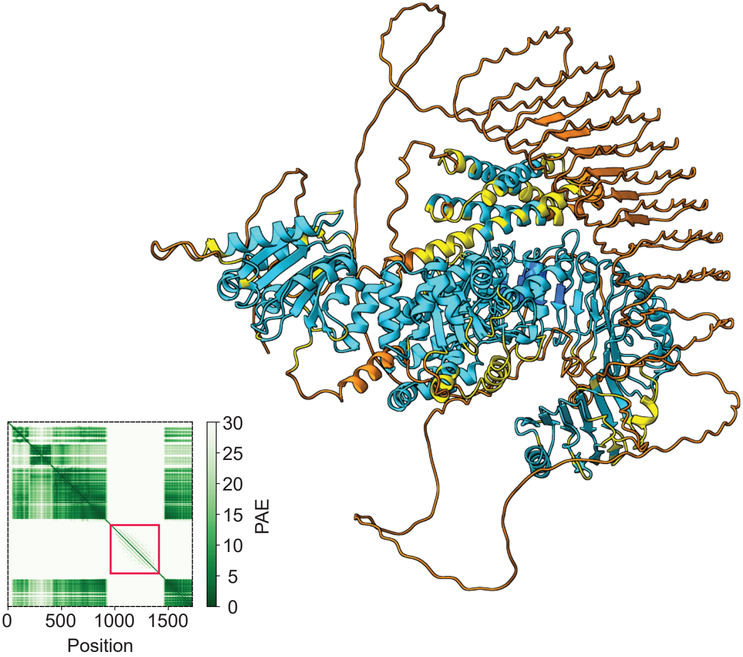


**Figure S20. Per-residue AlphaFold confidence and predicted aligned error plot for TrturKRN2B02G099510**

AlphaFold-predicted structure of TtrurKRN2B02G099510 colored by per-residue confidence (pLDDT): very high (blue), high (cyan), medium (yellow), and low (orange). The associated predicted aligned error (PAE) plot is shown on the left, with the region corresponding to simple repeats highlighted by a red box.

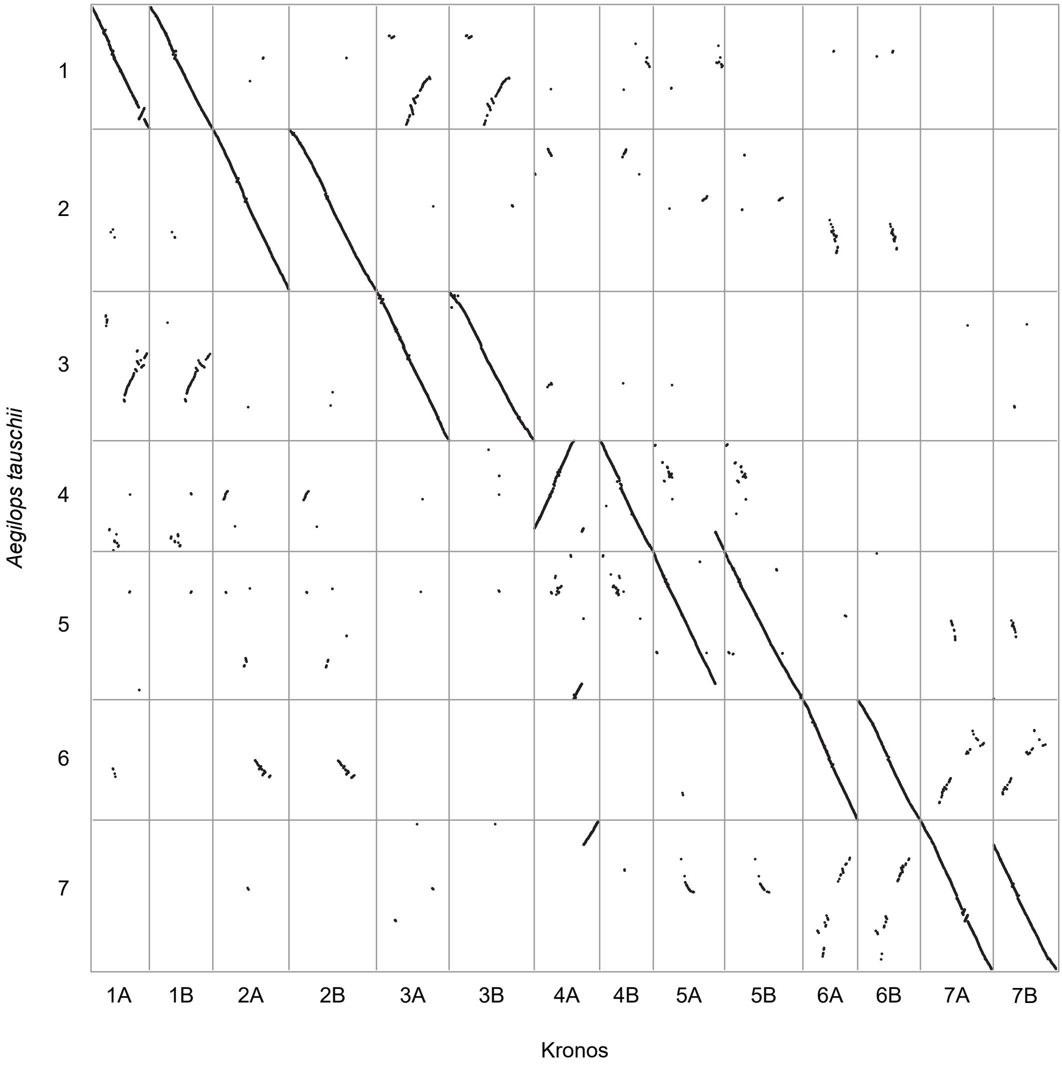


**Figure S21. Gene-level collinearity between Kronos and *Aegilops tauschii***

Dot plot displays syntenic relationships between annotated genes in Kronos and *Ae. tauschii* as identified by MCScanX. Each dot represents a pair of homologous genes detected within colinear blocks. Diagonal tracks indicate conserved chromosome-scale synteny, while off-diagonal signals reflect translocations, inversions, and lineage-specific rearrangements.

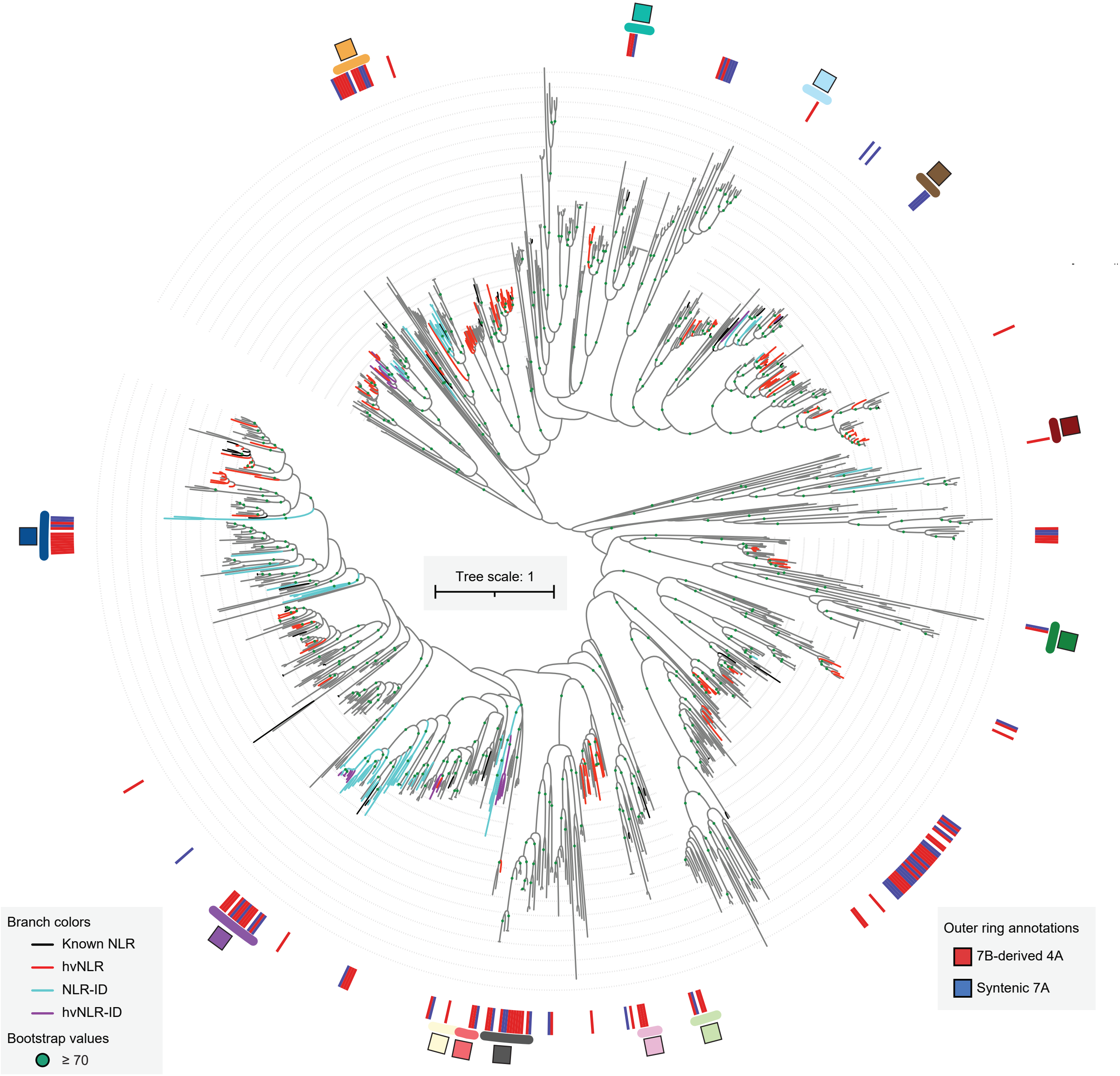


**Figure S22. Phylogenetic distribution of NLRs in the 7B-derived block of chromosome 4A and its collinear interval on chromosome 7A**

Maximum-likelihood phylogeny inferred from NB-ARC domains. Branch colors indicate: black—cloned, functionally validated NLRs; red—Kronos highly variable NLRs (hvNLRs); sky blue—Kronos NLR with PFAM-annotated integrated domains (NLR-ID); purple—Kronos hvNLR-IDs. All other Kronos NLRs are shown in gray. The outer ring denotes NLRs located in the 7B-derived block of chromosome 4A (red) and its colinear interval of chromosome 7A (blue). Colored arcs and boxes correspond to phylogenetic groups shown in Figure 4F.

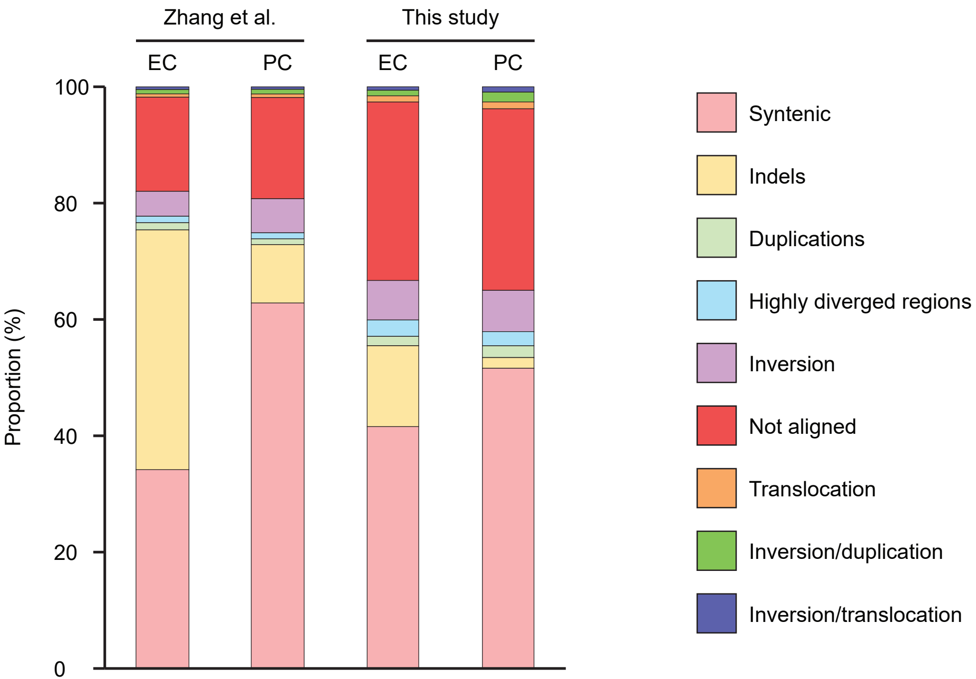


**Figure S23. Proportional distribution of EMS-induced mutations across genomic categories**

Uniquely identified mutations in Zhang et al. (2023) (left) and in this study (right) in exome-capture (EC) and promoter-capture (PC) datasets were assigned to each genomic category, identified by SyRi. The stacked bar plot displays the proportion of these mutations

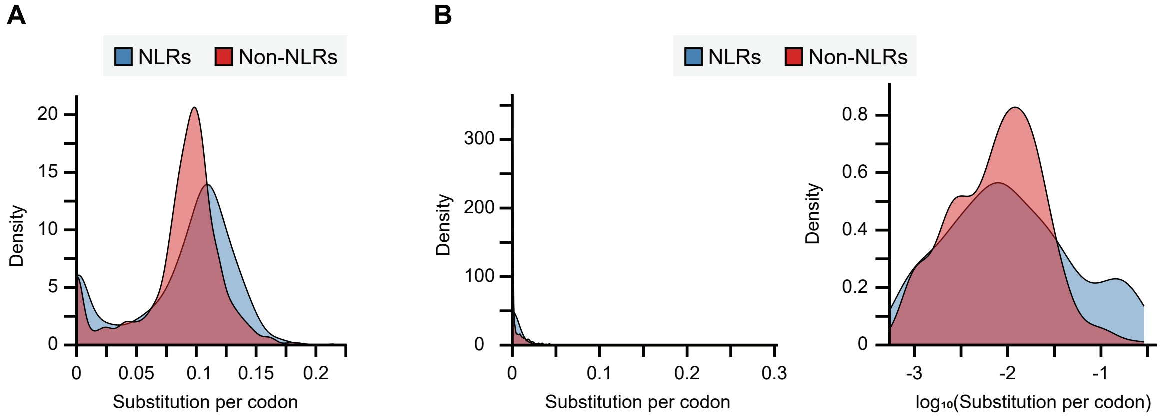


**Figure S24. Density of EMS-type substitutions per codon in NLRs versus non-NLRs**

Distributions of EMS-type mutations mapped to NLRs (blue) and non-NLR (red) genes are shown for **(A)** from uniquely mapped reads and **(B)** multi-mapped reads. For both panels, the number of substitutions was normalized by protein length (codons), resulting in the substitution rate per codon. Non-NLR genes were randomly sampled to match the length distribution of NLRs. **(B)** the $log_{10}$-transformed substitution rates (right) are displayed to improve visibility of the skewed distribution.

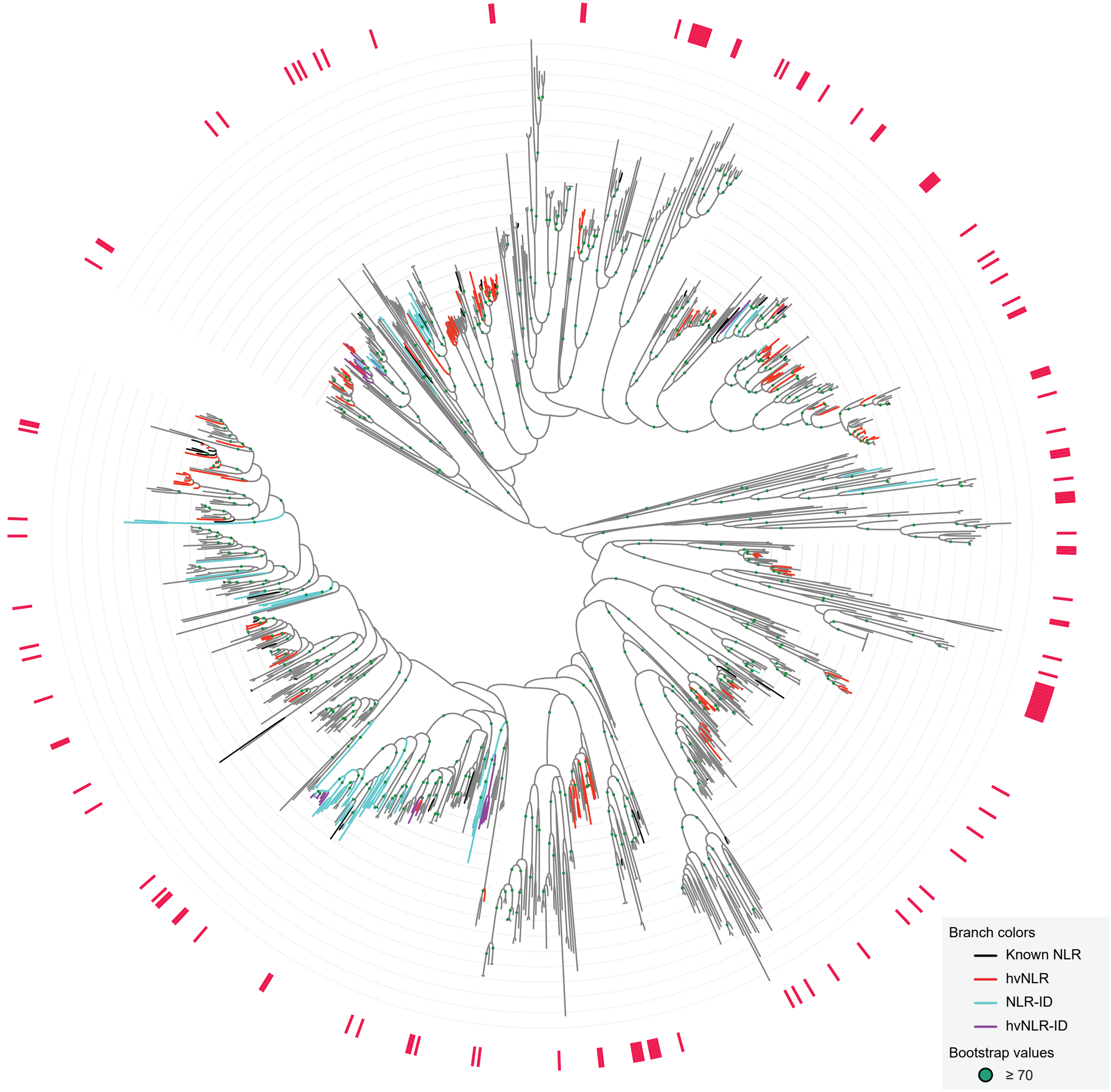


**Figure S25. Phylogenetic distribution of Kronos NLRs with low EMS mutation rates**

Maximum-likelihood phylogeny inferred from NB-ARC domains. Red boxes around the phylogeny indicates 135 NLRs that displayed < 0.03 EMS-type mutations per codon, based on exome-capture data. Branch colors indicate: black—cloned, functionally validated NLRs; red—Kronos highly variable NLRs (hvNLRs); sky blue—Kronos NLR with PFAM-annotated integrated domains (NLR-ID); purple—Kronos hvNLR-IDs. Bootstrap supports ≥ 70 are shown as green circles at the corresponding nodes.

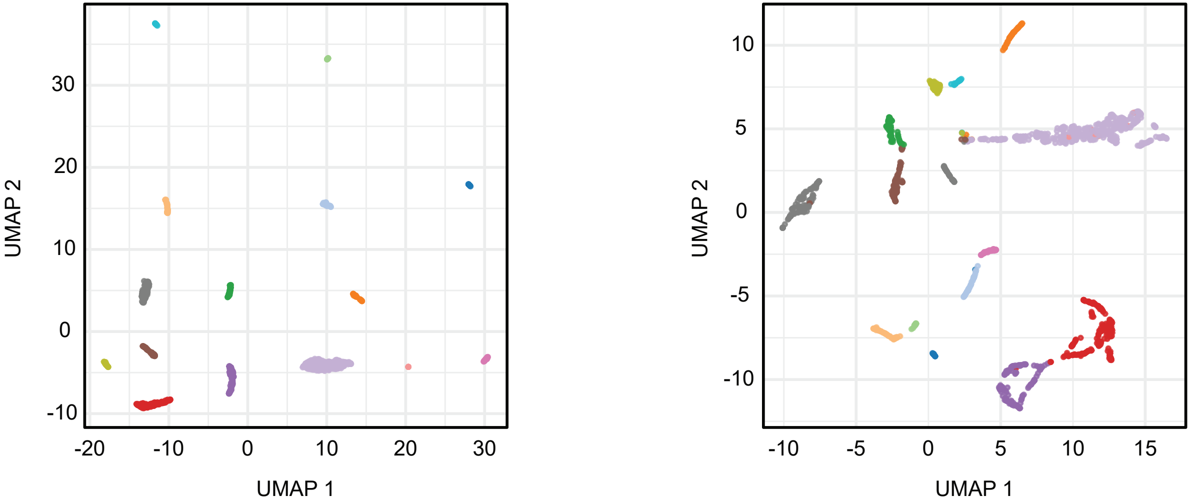


**Figure S26. Dimensionality reduction of EMS-type mutations using UMAP**

UMAP projections of EMS-type mutations are shown for exome-capture (left) and promoter-capture (right) datasets. Each point represents a Kronos mutant, and colors indicate cluster identities inferred from non-EMS-type mutations in the exome-capture dataset. These cluster assignments were applied consistently across both panels to facilitate direct comparison.

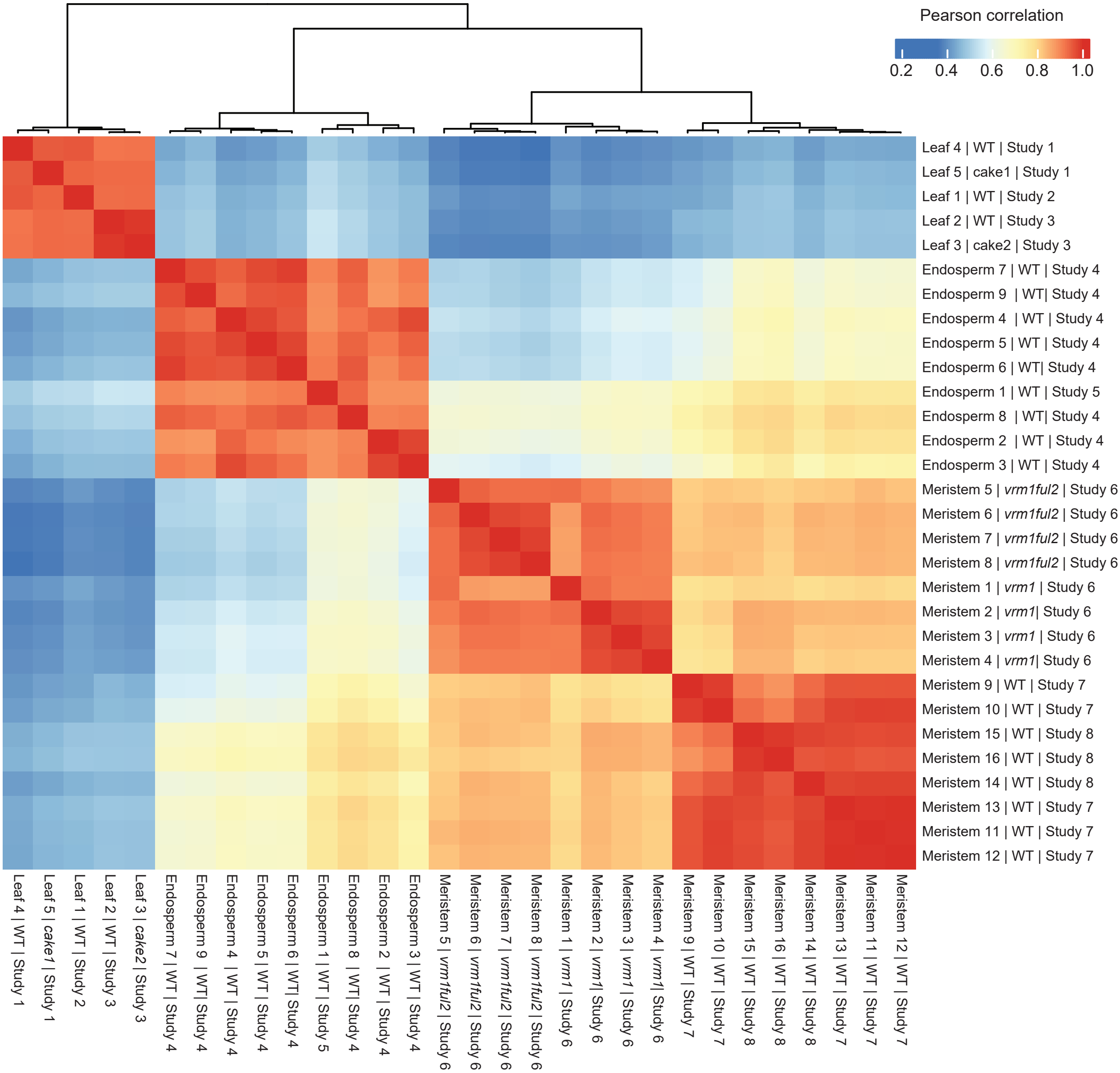


**Figure S27. Tissue-specific clustering of NLR expression profiles**

Heatmap of Pearson correlations between samples based on transcriptome-wide percentile ranks of NLRs. Samples are labeled by tissue, genotype, and study (Table S44). Hierarchical clustering highlights strong separation of leaf, endosperm, and meristem datasets across seven independent studies. Samples with distinct study numbers originate from different datasets.

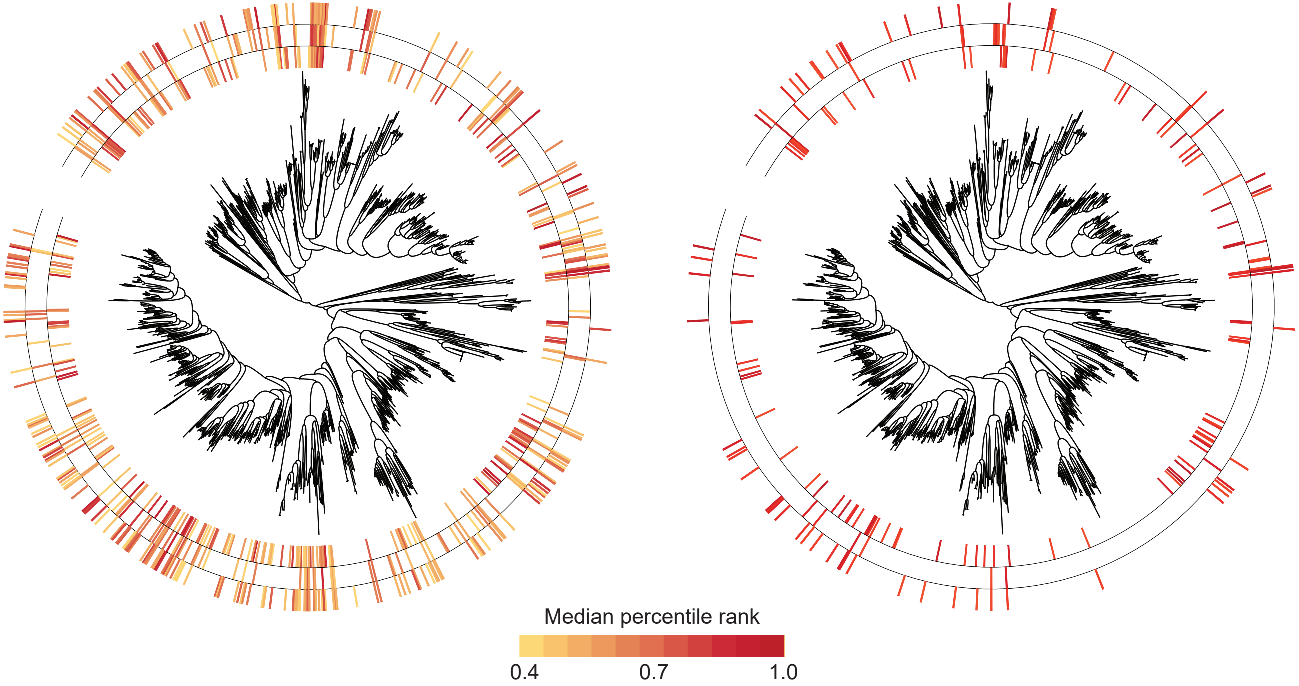


**Figure S28. Phylogenetic distributions of expressed NLRs across tissues**

Phylogenetic trees displaying NLRs with median percentile ranks ≥0.40 (left) and ≥0.70 (right) in leaf, endosperm, and meristem. Colored bars mapped onto the phylogeny indicate tissue-level median percentile ranks, with leaf (inner ring), endosperm (middle ring), and meristem (outer ring).

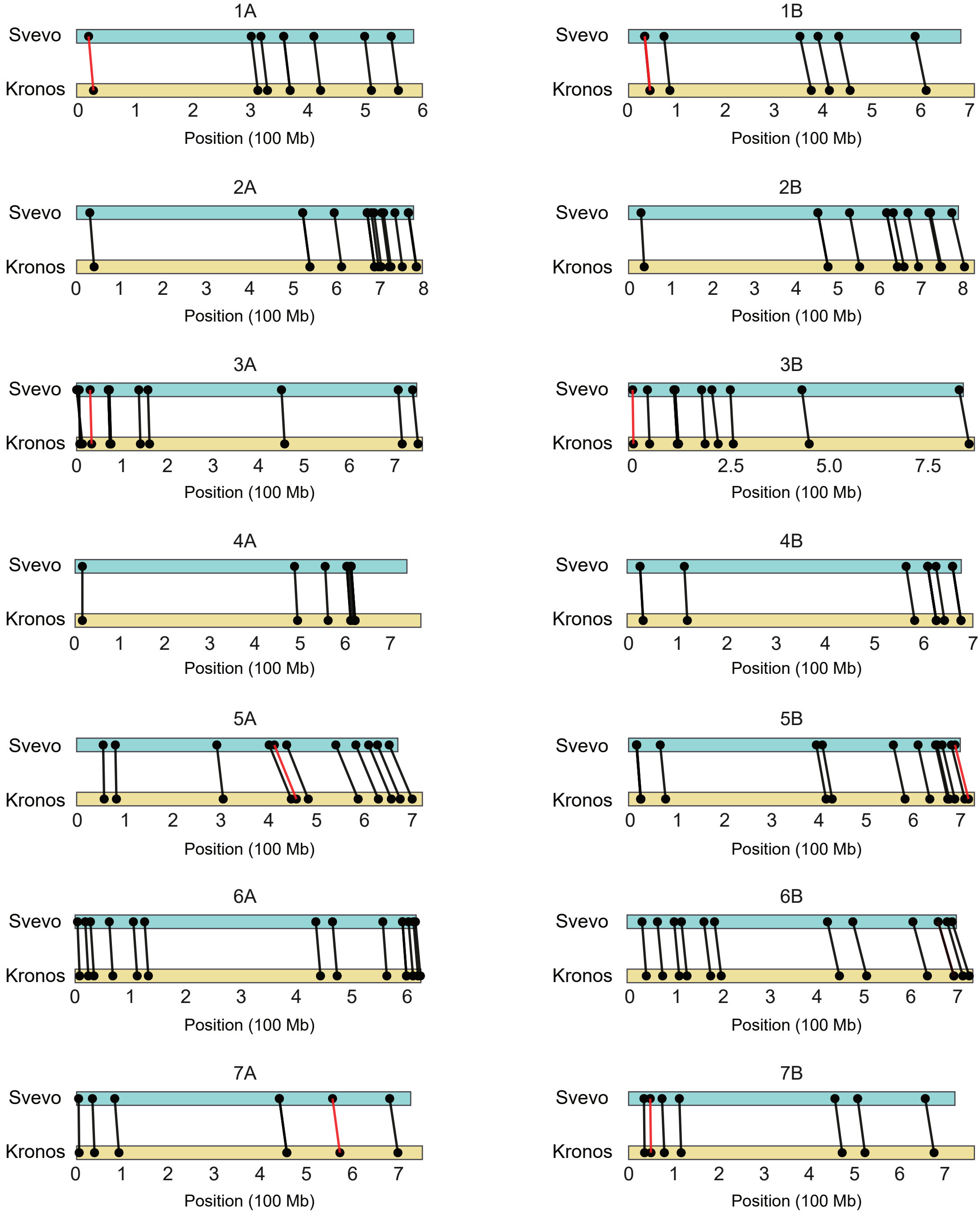


**Figure S29. Chromosome-wide distribution of miRNAs**

157 miRNA loci previously identified from Svevo were transferred to Kronos. Black lines indicate loci with 100% sequence identity, while red lines denote loci with minor sequence variation.

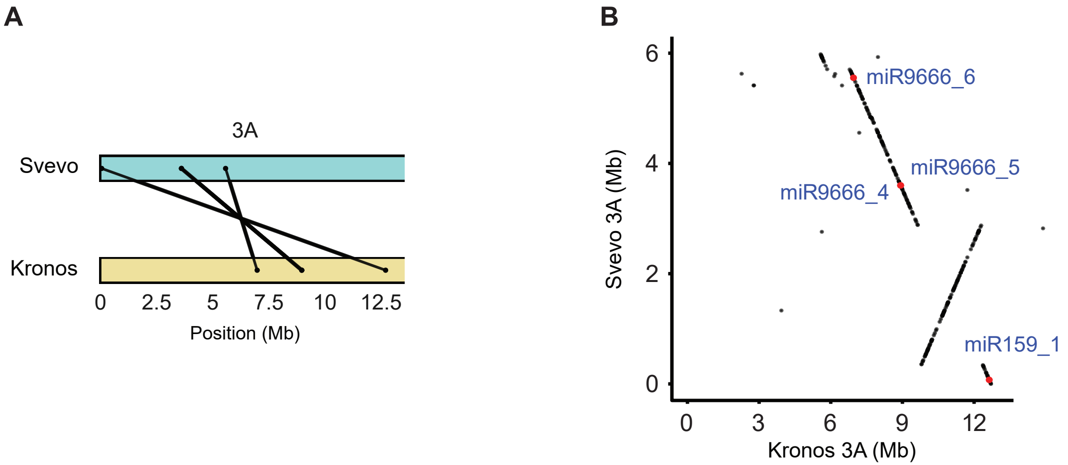


**Figure S30. Chromosome 3A inversion affecting miRNA positions between Svevo and Kronos**

**(A)** Positions of miRNA loci at the beginning of chromosome 3A (0–12 Mb in Kronos) are shown for Svevo and Kronos. **(B)** Local synteny plot highlighting an inversion in this region, which explains positional inconsistencies for miRNAs between Svevo and Kronos.

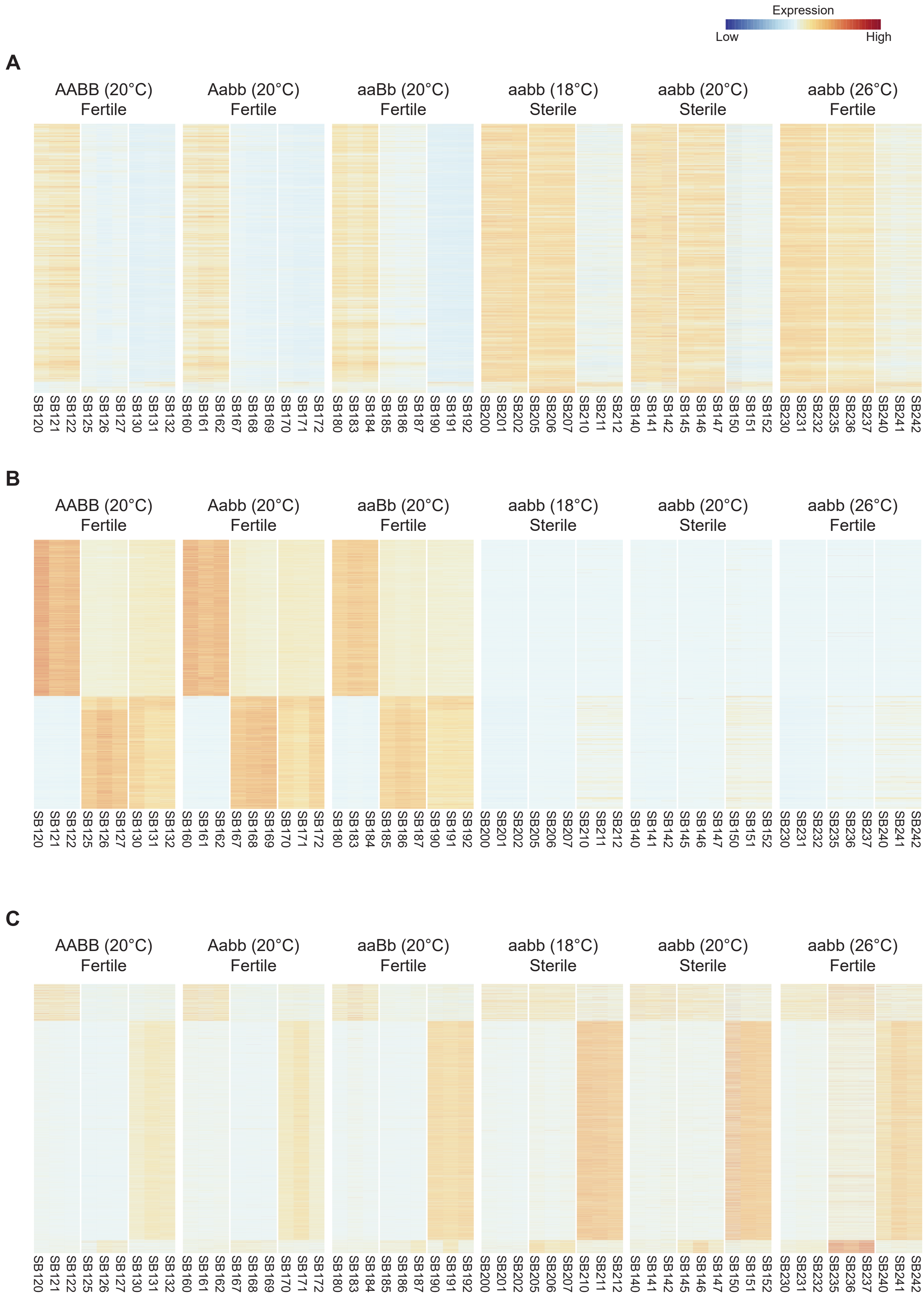


**Figure S31. Expression profiles of phasiRNAs and heterochromatic siRNAs across genotypes and conditions**

Heatmaps showing expression profiles of small RNAs across sRNA transcriptome datasets (Table S49). **(A)** 21-nt phasiRNAs. **(B)** 24-nt phasiRNAs. **(C)** 24-nt heterochromatic siRNAs (hc-siRNAs). Samples are grouped by genotype (AABB, Aabb, aaBb, aabb) and condition (temperature and fertility status). Expression values are scaled from low (blue) to high (red). Sample names along the x-axis match those listed in Table S49.

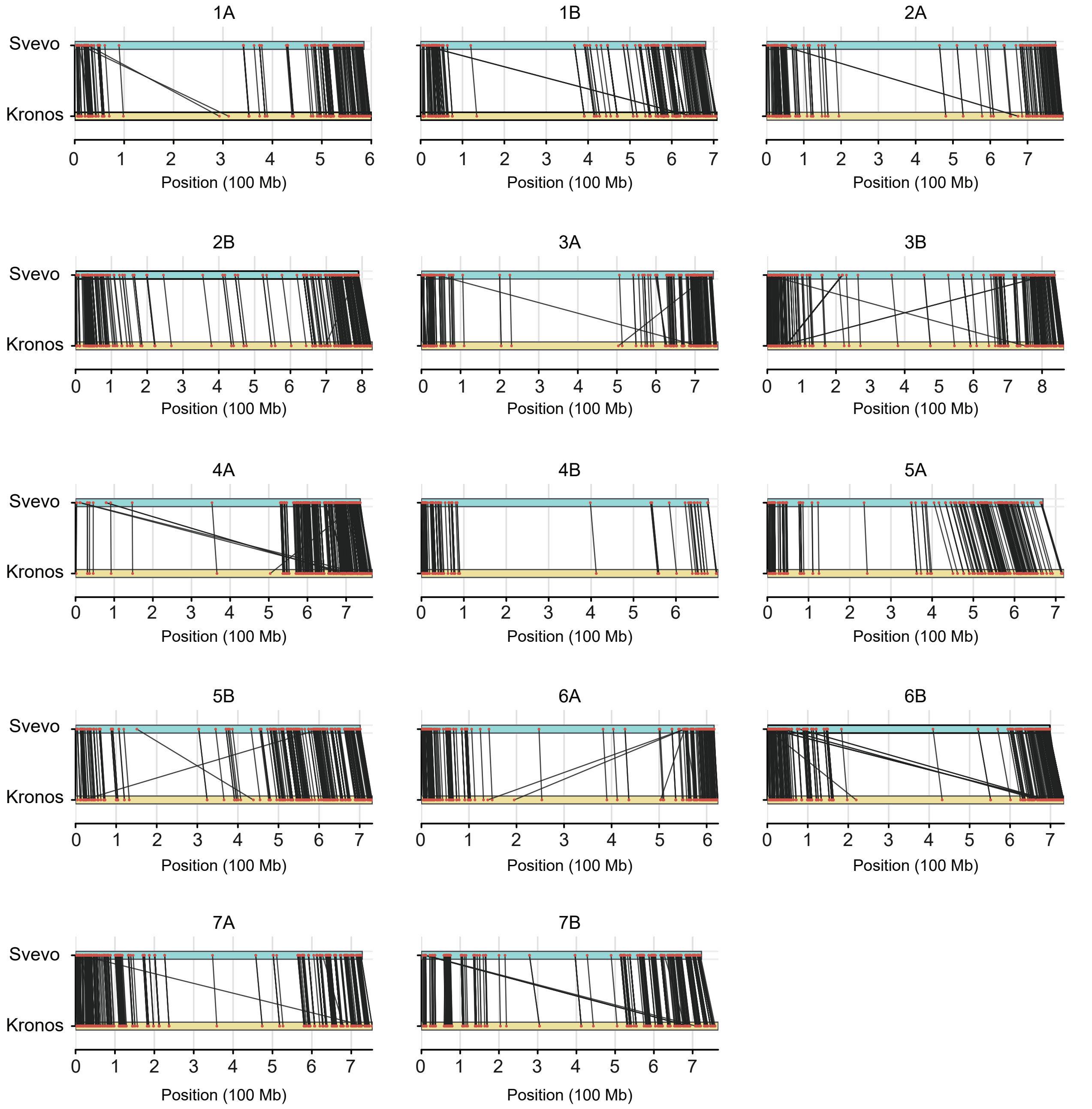


**Figure S32. Chromosome-scale collinearity of *PHAS* loci between Svevo and Kronos**

Reciprocal similarity searches of 9,104 *PHAS* loci annotated in Svevo and mapped to the Kronos genome. Each panel shows alignments of *PHAS* loci between corresponding chromosomes of Svevo (top) and Kronos (bottom). Black lines indicate syntenic placements of loci, with top hits plotted on syntenic chromosomes.

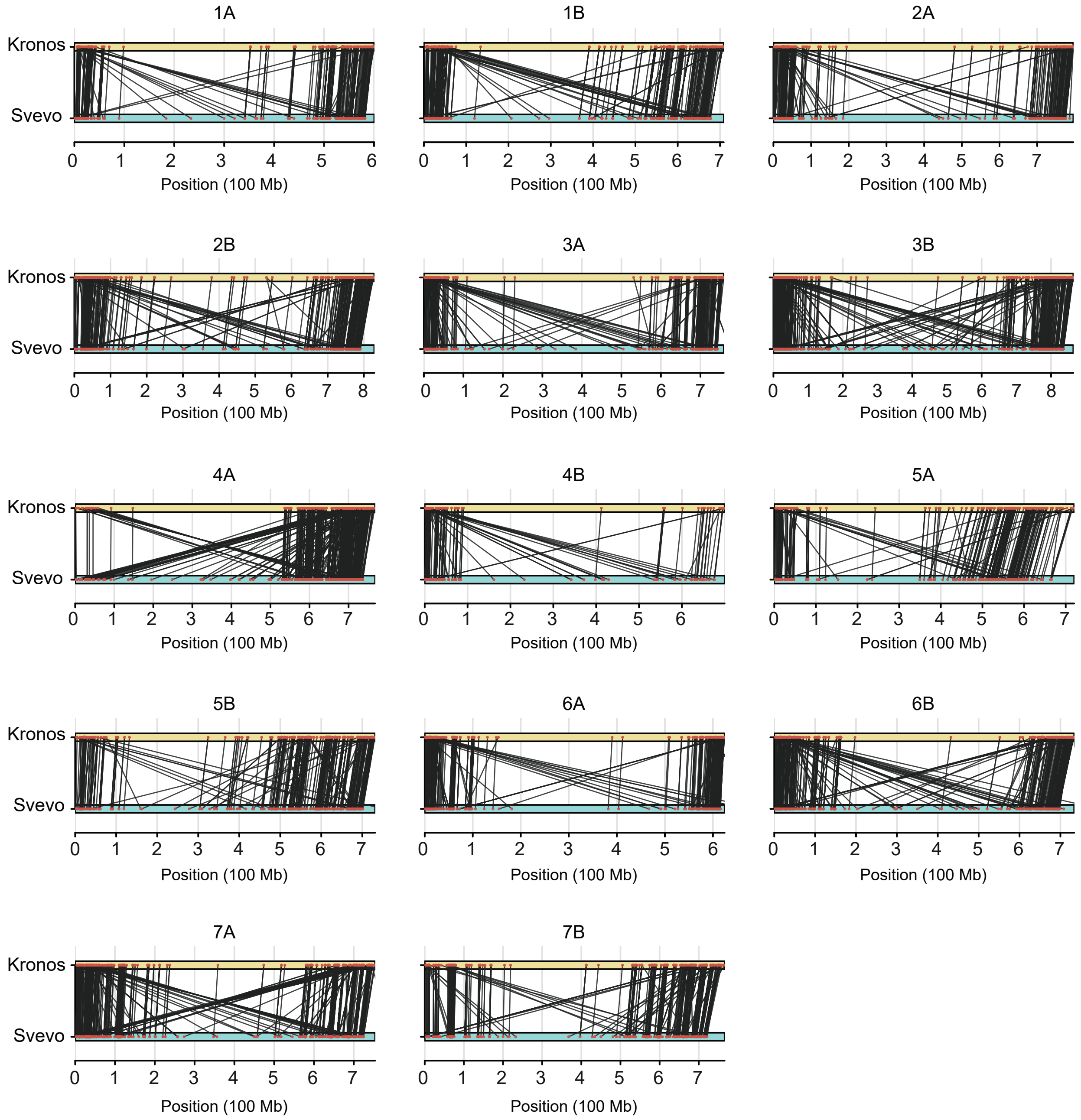


**Figure S33. Chromosome-scale collinearity of *PHAS* loci between Kronos and Svevo**

Reciprocal similarity searches of 10,965 *PHAS* loci annotated in Kronos and mapped to the Svevo genome. Each panel shows alignments of *PHAS* loci between corresponding chromosomes of Kronos (top) and Svevo (bottom). Black lines indicate syntenic placements of loci, with top hits plotted on syntenic chromosomes.
